# The development of aperiodic neural activity in the human brain

**DOI:** 10.1101/2024.11.08.622714

**Authors:** Zachariah R. Cross, Samantha M. Gray, Adam J. O. Dede, Yessenia M. Rivera, Qin Yin, Parisa Vahidi, Elias M. B. Rau, Christopher Cyr, Ania M. Holubecki, Eishi Asano, Jack J. Lin, Olivia Kim McManus, Shifteh Sattar, Ignacio Saez, Fady Girgis, David King-Stephens, Peter B. Weber, Kenneth D. Laxer, Stephan U. Schuele, Joshua M. Rosenow, Joyce Y. Wu, Sandi K. Lam, Jeffrey S. Raskin, Edward F. Chang, Ammar Shaikhouni, Peter Brunner, Jarod L. Roland, Rodrigo M. Braga, Robert T. Knight, Noa Ofen, Elizabeth L. Johnson

## Abstract

The neurophysiological mechanisms supporting brain maturation are fundamental to attention and memory capacity across the lifespan. Human brain regions develop at different rates, with many regions developing into the third and fourth decades of life. Here, in this preregistered study (https://osf.io/gsru7), we analyzed intracranial EEG (iEEG) recordings from widespread brain regions in a large developmental cohort. Using task-based (i.e., attention to-be-remembered visual stimuli) and task-free (resting-state) data from 101 children and adults (5.93 – 54.00 years, 63 males; *n* electrodes = 5691), we mapped aperiodic (1/ƒ-like) activity, a proxy of neural noise, with steeper slopes indexing less noise and flatter slopes indexing more noise. We reveal that aperiodic slopes flatten with age into young adulthood in both association and sensorimotor cortices, challenging models of early sensorimotor development based on brain structure. In prefrontal cortex (PFC), attentional state modulated age effects, revealing steeper task-based than task-free slopes in adults and the opposite in children, consistent with the development of cognitive control. Age-related differences in task-based slopes also explained age-related gains in memory performance, linking the development of PFC cognitive control to the development of memory. Last, with additional structural imaging measures, we reveal that age-related differences in gray matter volume are similarly associated with aperiodic slopes in association and sensorimotor cortices. Our findings establish developmental trajectories of aperiodic activity in localized brain regions and illuminate the development of PFC control during adolescence in the development of attention and memory.

## Introduction

Human brain regions develop at different rates, with many regions developing into the third and fourth decades of life, followed by gradual declines in volume throughout adulthood (Bethlehem et al., 2022; Gogtay et al., 2004; Grydeland et al., 2019). Understanding the complexities of human brain development requires a comprehensive investigation into the intricate interplay between electrophysiological dynamics, brain structure, and behavior across the lifespan. Despite the importance of this endeavor to basic and translational neuroscience, research has been limited by a paucity of methods capable of studying human brain function with high spatial and temporal precision and focused on narrow age ranges. Further, non-oscillatory, aperiodic activity has yet to be fully characterized from a developmental perspective (cf. (Favaro et al., 2023; A. T. Hill et al., 2022; Schaworonkow & Voytek, 2021; Tröndle et al., 2022). Consequently, the manifestation of age-related differences in aperiodic activity and their relation to brain structure and cognition remain unknown. The aperiodic component of the electrophysiological power spectrum, characterized by its spectral slope and offset (Donoghue et al., 2020; Wen & Liu, 2016), is hypothesized to reflect neural noise (Ahmad et al., 2022; van Nifterick et al., 2023), with a flatter slope and lower offset posited to reflect increased excitatory neuronal population spiking (Manning et al., 2009; K. J. Miller et al., 2012). While there are several interpretations of the aperiodic signal, it is likely produced by a variety of biological mechanisms (Kramer & Chu, 2024; Voytek et al., 2015; Voytek & Knight, 2015), including low-pass filtering property of dendrites (Buzsáki et al., 2012), frequency dependence of current propagation in biological tissues (Bédard & Destexhe, 2009), stochastically driven damped oscillators with different relaxation rates (Evertz et al., 2022), and/or the balance between excitation and inhibition (E/I) in neuronal populations (Martínez-Cañada et al., 2023; Wiest et al., 2023). From this perspective, these broad range of biological factors likely make up the aperiodic signal and reflect age-related differences in neural ‘noise’.

The balance of an optimal level of neural noise is a fundamental property of healthy brain function (Turrigiano & Nelson, 2004). Indeed, an optimal level neural noise is proposed to safeguard against hyper-synchronization, with imbalances in noise implicated in neurodevelopmental disorders, such as schizophrenia and autism (Earl et al., 2024; Pani et al., 2022; Shuffrey et al., 2022) and generalized learning disabilities (Fernandez & Garner, 2007). Studies using scalp electroencephalography (EEG) during passive (i.e., task-free) states have consistently demonstrated a flattening of the slope and a downward shift in the offset with advancing age throughout adulthood (Donoghue et al., 2020; Merkin et al., 2023; Voytek et al., 2015; Waschke et al., 2017). Such age-related flattening in task-free aperiodic activity predicts declines in memory performance (Voytek et al., 2015) and alterations in stimulus-related neurophysiological responses, such as inter-trial alpha phase clustering during visual spatial discrimination in the elderly (Tran et al., 2020). By contrast, flatter task-based aperiodic slopes are associated with enhanced memory and learning in healthy young adults (Cross et al., 2022; Lendner et al., 2023), hinting at a nuanced interplay between aperiodic activity, attentional state, and age. Thus, understanding the development of aperiodic activity and its modulation by attentional state, with high spatial precision, is necessary to understand brain development and cognitive function across the lifespan.

To date, developmental studies of aperiodic activity have relied on scalp EEG (Cellier et al., 2021; Favaro et al., 2023; A. T. Hill et al., 2022; Schaworonkow & Voytek, 2021). Yet, scalp-EEG is limited in spatial resolution and cannot reliably characterize regionally precise neurophysiological activity (Ofen et al., 2019). To overcome these limitations, we analyzed rare intracranial EEG (iEEG) data from an exceptionally large developmental cohort of neurosurgical patients aged 5 to 54 years undergoing invasive monitoring for seizure management. In contrast to noninvasive neuroimaging, iEEG provides both spatially localized information and the high temporal precision needed to examine neurophysiology (Johnson et al., 2020; Johnson & Knight, 2015; Parvizi & Kastner, 2018), and is thus an invaluable tool for investigating mechanisms of cognitive and brain maturation (Johnson, Tang, et al., 2018; Johnson, Yin, et al., 2022; Johnson & Knight, 2023; Ofen et al., 2019; Rau et al., n.d.; Yin et al., 2020, 2023). iEEG provides rich and novel measures of neurophysiology including low-frequency periodic and aperiodic activity, and high-frequency broadband activity reflecting neuronal population activity (Leszczyński et al., 2020; Nir et al., 2007; Ray et al., 2008; Rich & Wallis, 2017; Watson et al., 2018). Thus, iEEG enables unique discoveries of the neurophysiological mechanisms of cognitive and brain maturation in humans.

In this preregistered study (https://osf.io/gsru7), we sought to define regionally precise, brain-wide developmental trajectories of aperiodic activity in task-based and task-free states (Figures 1A, 1B). In addition to mapping aperiodic activity across development, we defined the relationship between regionally precise aperiodic activity and cortical structure (Figure 1C). Measures of regional gray matter volume (GMV) and electrophysiological activity show substantial overlap in relation to cognition, pathology (Hunt et al., 2016; Schölvinck et al., 2013), and age (Doval et al., 2024; Overbye et al., 2018; Sui et al., 2014; Whitford et al., 2007), which suggests that they may be jointly explained by shared factors, such as myelination and synaptogenesis. Thus, examining structure-function coupling can provide context to understand novel electrophysiological findings, such as iEEG measures of aperiodic activity by age, based on well-documented age-related variability in regional brain structure (Bethlehem et al., 2022; Gogtay et al., 2004; Groeschel et al., 2010). Based on reports of age-related variability in global scalp EEG-derived aperiodic activity (Cellier et al., 2021; Finley et al., 2022; A. T. Hill et al., 2022; Schaworonkow & Voytek, 2021; Thuwal et al., 2021) and in brain structure demonstrating that sensorimotor regions mature earlier than association regions (Gogtay et al., 2004; Grydeland et al., 2019; J. Hill et al., 2010; Sydnor et al., 2021), we hypothesized that: (a) in association cortices, the aperiodic slope flattens with age into young adulthood; (b) in sensorimotor cortices, the aperiodic slope flattens with age into adolescence; (c) attentional state (task-based vs. task-free) modulates age effects observed in (a) and (b), and; (d) age-related differences in aperiodic activity are modulated by regional GMV.

**Figure 1.**
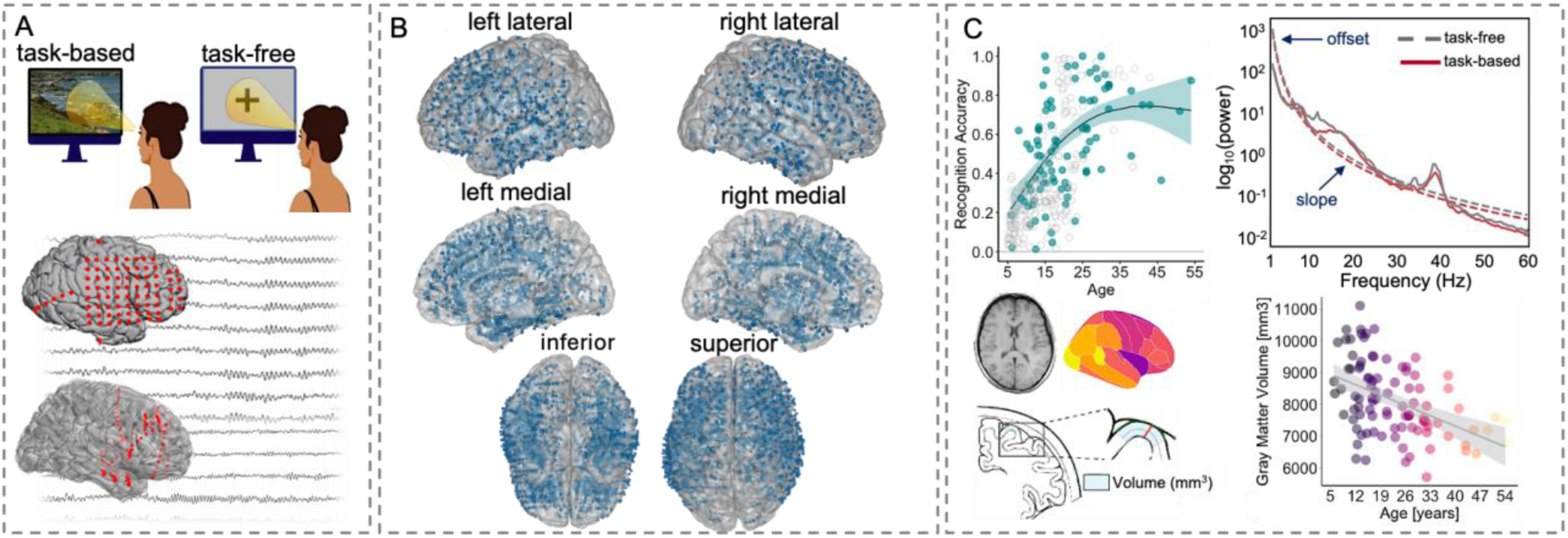
Design, channel coverage, and key variables. **(A)** Intracranial neurophysiological activity was recorded using electrocorticography (ECoG; middle) and stereoelectroencephalography (sEEG; bottom) during both task-based (top left) and task-free wake states (top right). **(B)** Seizure- and artifact-free intracranial channel placements (*n* = 5691) across all patients (*n* = 101) in MNI space. **(C)** Schematic of key dependent and independent variables. Top left: iEEG patients (teal; *n* = 81) show the expected developmental trajectory of improved memory recognition from ∼5 – 30 years of age (*p* ≤.001) and fall in the range of age-matched, healthy controls (gray; *n* = 221). Top right: schematic power spectral density plot illustrating the periodic (oscillatory) components over and above the aperiodic (1/ƒ-like) component in task-free (dashed) and task-based (solid) conditions. The offset (i.e., y-intercept) and slope (exponent) make up the aperiodic component when power (y-axis) is in log-space. Bottom left: TI MRI obtained for each patient, parcellation of cortical regions based on the Desikan-Killiany-Tourville atlas, and GMV estimation (adapted from Bethlehem et al., 2022). Bottom right: age-related differences in global GMV (mm^3^) in our cohort, showing the expected developmental trajectory of decreased GMV from ∼5 – 54 years of age (*p* ≤.001).

We first reveal a gradient in aperiodic activity across the brain, suggesting less neural noise in inferior lateral posterior regions and more neural noise in superior medial frontal regions. We then establish developmental trajectories of aperiodic activity, revealing a flattening of the slope from childhood to young adulthood in both association and sensorimotor cortices, challenging our hypothesis that aperiodic activity stabilizes before young adulthood in sensorimotor cortices. We further reveal how attentional state modulates age effects in select regions including prefrontal cortex (PFC) and establish predictive links between task-based aperiodic activity in PFC and individual memory outcomes. We last uncover novel associations between cortical structure and function, highlighting how aperiodic slopes in both association and sensorimotor cortices are modulated by age-related variability in GMV. Taken together, we offer critical insights into the intricate interplay between aperiodic neural activity, cortical structure, and behavior from childhood to middle age and illuminate the development of PFC control during adolescence in the development of attention and memory.

## Results

### iEEG memory and brain volume measures generalize to healthy populations

One hundred and one neurosurgical patients participated (mean age = 19.25, range = 5.93 – 54.00 years; 63 males). Patients were selected based on above-chance behavioral performance on two visual memory recognition tasks (mean normalized accuracy = 0.54, SD = 0.25, range = 0.01 – 1.00; *β* = 0.54, SE = 0.02, *p* ≤.001) and/or if there was a task-free recording available. Those with major lesions, prior surgical resections, noted developmental delays, or neuropsychological memory test scores <80 were considered ineligible. A nonlinear regression (single spline with two internal knots) of recognition accuracy by age indicated a positive association (first knot: *β* = 0.90, SE = 0.16, *p* ≤.001; second knot: *β* = 0.30, SE = 0.13, *p* = .02; *R^2^* = .27; see Figure 1C), indicating that iEEG patients exhibit the expected developmental trajectory of improved memory from age 5-30 years, consistent with age-matched, healthy controls (Johnson, Yin, et al., 2022; Ofen et al., 2007, 2019). Analysis of global GMV by age indicated a negative association (*β* = -46.56, SE = 9.36, *p* ≤.001, *R^2^* = .20; Figure 1C), indicating that with every one-year increase in age there is a 46mm^3^ reduction in GMV. This further demonstrates that iEEG patients show the expected developmental trajectory of decreased GMV from age 5 to 54 years, consistent with well-documented decreases in GMV from childhood through adulthood in healthy individuals (Bethlehem et al., 2022; Gogtay et al., 2004; Groeschel et al., 2010; Wilke et al., 2007). These demonstrations provide converging evidence that the results of our iEEG analyses generalize to healthy populations (P. F. Hill et al., 2020; Johnson & Knight, 2023).

### Aperiodic activity differs by brain region

Prior to testing hypotheses, we first characterized regional differences in aperiodic activity by implementing linear mixed-effects models, regressing region onto the aperiodic slope while regressing out attentional state (task-based, task-free) and age, treating participants and nested channels as random intercepts (Johnson & Knight, 2023). Regions of interest (ROI) were defined based on the Desikan-Killiany-Tourville (DKT) atlas (Klein & Tourville, 2012). The goodness-of-fit (i.e., *R^2^*) between task-based (*M* = 0.93, *SD* = 0.05) and task-free (*M* = 0.94, *SD* = 0.05) conditions indicated good fits (see Fig. S15 for the distribution of model fits between conditions). We revealed a gradient of steeper slopes in inferior lateral posterior regions to flatter slopes in superior medial frontal regions (χ2(19) = 1038.30, *p* ≤ 0.001; Figure 2A). These results extend previous reports of a posterior-to-anterior gradient in task-free neural noise based on fMRI, i.e., Hurst exponent (Fotiadis et al., 2023) and magnetoencephalography (MEG) aperiodic component (Mahjoory et al., 2020). Our data demonstrate that aperiodic activity varies between localized brain regions to higher degree than previously reported.

**Figure 2.**
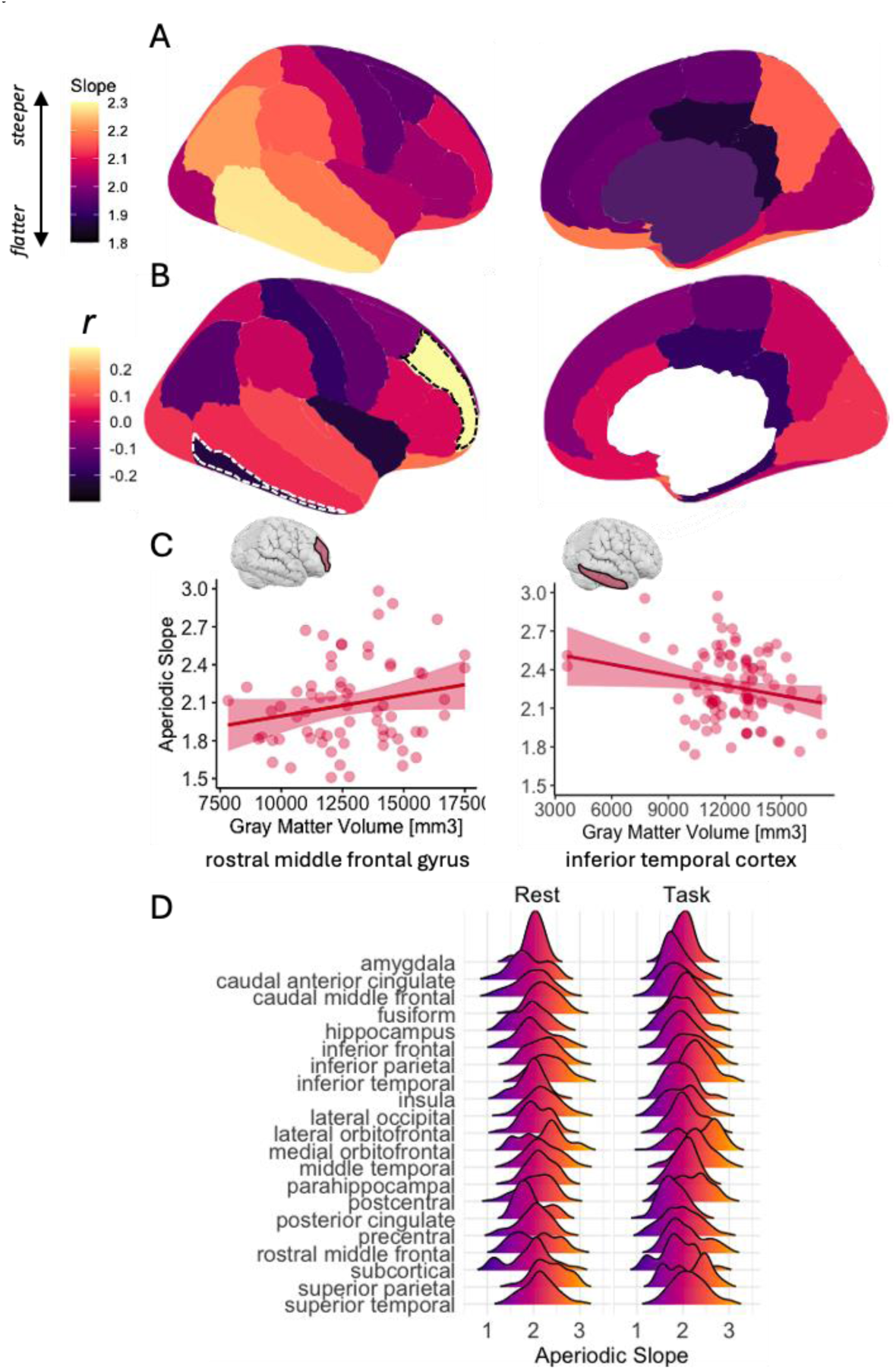
Regional differences in the aperiodic slope and correlations with GMV. **(A)** Brain-wide standardized means (predicted marginal means) of regional aperiodic slopes adjusted across attentional state, age and the random effects structures. Warmer colors/higher values indicate steeper slopes. (**B)** Brain-wide correlations (Spearman rho) between regional GMV (mm^3^) and aperiodic slopes. Warmer colors/higher values indicate positive correlations; cooler colors/lower values indicate negative correlations. Note that the area corresponding to subcortical space is white as no analysis of subcortical GMV was performed. Regions with statistically significant correlations (*p* < 0.05) are indicated by dashed borders. **(C)** Scatterplots illustrating relationships between GMV (x-axis) and aperiodic slopes in regions with statistically significant correlations. Individual data points represent single participant data averaged across channels for each representative ROI. Shading shows the standard error. (**D**) Ridgeline plot illustrating the distribution of aperiodic slopes (x-axis; higher values denote a steeper slope) by region (y-axis) and condition (left: task-free; right: task-based).

Second, to characterize relationships between regional GMV and aperiodic activity (i.e., structure-function coupling), we correlated regional aperiodic slopes with regional GMV. We revealed regionally specific relationships between aperiodic activity and GMV (Figure 2C and Figure 2D). We observed a significant positive correlation with slope and GMV in rostral middle frontal gyrus (*r* = 0.27, *p*_adj_ = 0.03, 95% CI = [0.04, 0.48]), and a significant negative correlation in inferior temporal cortex (*r* = -0.22, *p*_adj_ = 0.03, 95% CI = [-0.40, -0.01]). These results reveal opposing structure-function coupling between localized regions and indicate that there is not a one-to-one mapping between GMV and aperiodic activity.

### Aperiodic activity stabilizes in young adulthood in associative and sensorimotor cortices

Having demonstrated that aperiodic activity differs by brain region, we sought to establish developmental trajectories of aperiodic activity between association and sensorimotor cortices. We first examined hypotheses a, that in association cortices, the aperiodic slope flattens with age into young adulthood, and b, that in sensorimotor cortices, the aperiodic slope flattens with age into adolescence (see Table S1 for a summary of association and sensorimotor regions). We implemented nonlinear mixed-effects regressions, modeling aperiodic activity as a function of age (fit with one spline; two knots) and cortex type (association, sensorimotor; Table S1), treating participant and DKT region as random effects on the intercept, and channel nested under participant. We revealed a significant age × cortex type interaction (*β* = 0.22 [95% CI = 0.14, 0.30], SE = 0.04, *p* < 0.001; age: *β* = -0.64 [95% CI = -0.99, -0.30], SE = 0.17, *p* < 0.001; cortex type: *β* = -0.04 [95% CI = -0.12, 0.03), SE = 0.04, *p* = 0.31; see Figure S1 for model diagnostics), demonstrating that the slope flattens with age into young adulthood, with the greatest flattening difference in sensorimotor compared to association cortices at 30 years of age (*β* = -0.19, *SE* = 0.08, *p* = 0.02; Figure 3). These results support our hypothesis that the aperiodic slope flattens with age into young adulthood in association cortices. These results are contrary to our hypothesis that aperiodic activity stabilizes with age into adolescence in sensorimotor cortices; however, the difference in flattening suggests dissociable trajectories between association and sensorimotor cortices.

**Figure 3.**
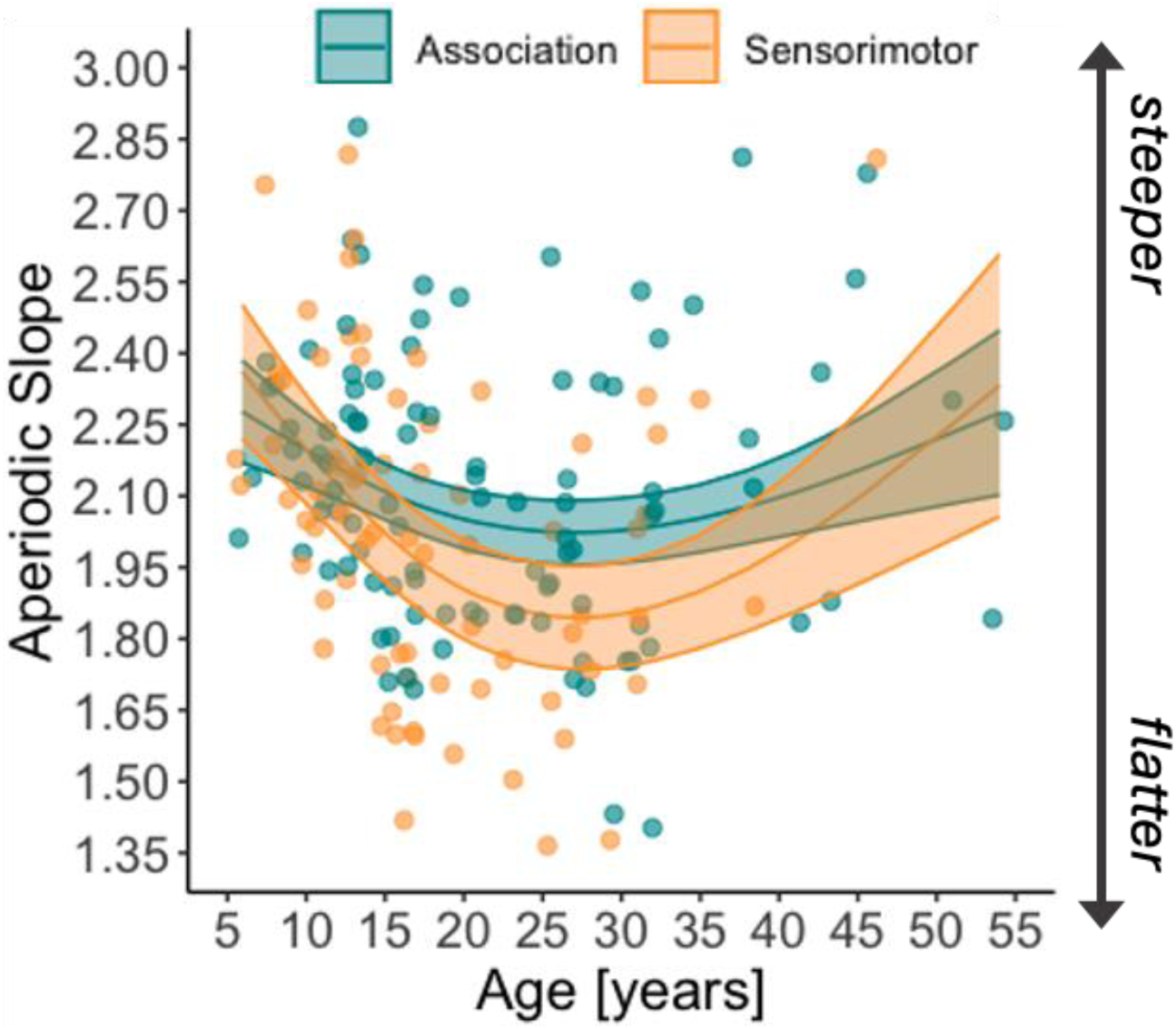
Age-related differences in aperiodic slopes between association and sensorimotor cortices. Modelled effects for differences in the aperiodic slope (y-axis; higher values denote a steeper slope) by age (x-axis). Association cortices are presented in teal and sensorimotor cortices in orange. Shaded regions indicate the 83% confidence interval. Individual data points represent slope values per participant averaged over channels.

### Regional aperiodic neural activity differs by age and attentional state

We next sought to establish developmental trajectories of aperiodic activity within localized brain regions and test hypothesis c, that age effects would differ between attentional states. To identify regional age effects in aperiodic activity and whether they differ by attentional state, we implemented separate linear mixed-effects models for each ROI. Our strategy for each analysis was to fit a model to the aperiodic slope and regress the estimates onto age (in years), attentional state (task-based, task-free), and the interaction of age and attentional state. All models were fit with by-participant and by-task random intercepts, with channel nested under participant (see Figures S2 and S3 for model diagnostics).

We revealed significant age × attentional state interactions in caudal middle frontal gyrus (cMFG; *β* = -0.003 [95% CI = -0.006, -0.002], SE = 0.001, *p*_adj_ = 0.007; age: *β* = -0.005 [95% CI = - 0.01, -0.003], SE = 0.004, *p*_adj_ = 0.41; task: *β* = 0.09 [95% CI = 0.05, 0.13], SE = 0.02, *p*_adj_ ≤ 0.001) and rostral middle frontal gyrus (rMFG; *β* = -0.004 [95% CI = -0.008, -0.002], SE = 0.001, *p*_adj_ = 0.04; age: *β* = -0.01 [95% CI = -0.03, -0.007], SE = 0.004, *p*_adj_ = 0.02; task: *β* = 0.09 [95% CI = 0.02, 0.15], SE = 0.03, *p*_adj_ = 0.37). In both cMFG and rMFG, task-free slopes are steeper than task-based slopes in children and the opposite is observed in adults; the direction of differences reverses around age 18 – 20 years (Figure 4B). If flatter slopes imply greater neural noise, and PFC activity reflects activation of cognitive control, then these results are consistent with increased cognitive control during task engagement in adolescence (Keller et al., 2023; Larsen et al., 2023; Sydnor et al., 2021; Sydnor et al., 2023) and mirror the development of domain-general cognitive control (Tervo-Clemmens et al., 2023). These results also support our hypothesis that attentional state modulates age-related flattening of the aperiodic slope. For visualizations of the main effects of age and condition on the aperiodic slope, see Figures S4 and S5, respectively.

**Figure 4.**
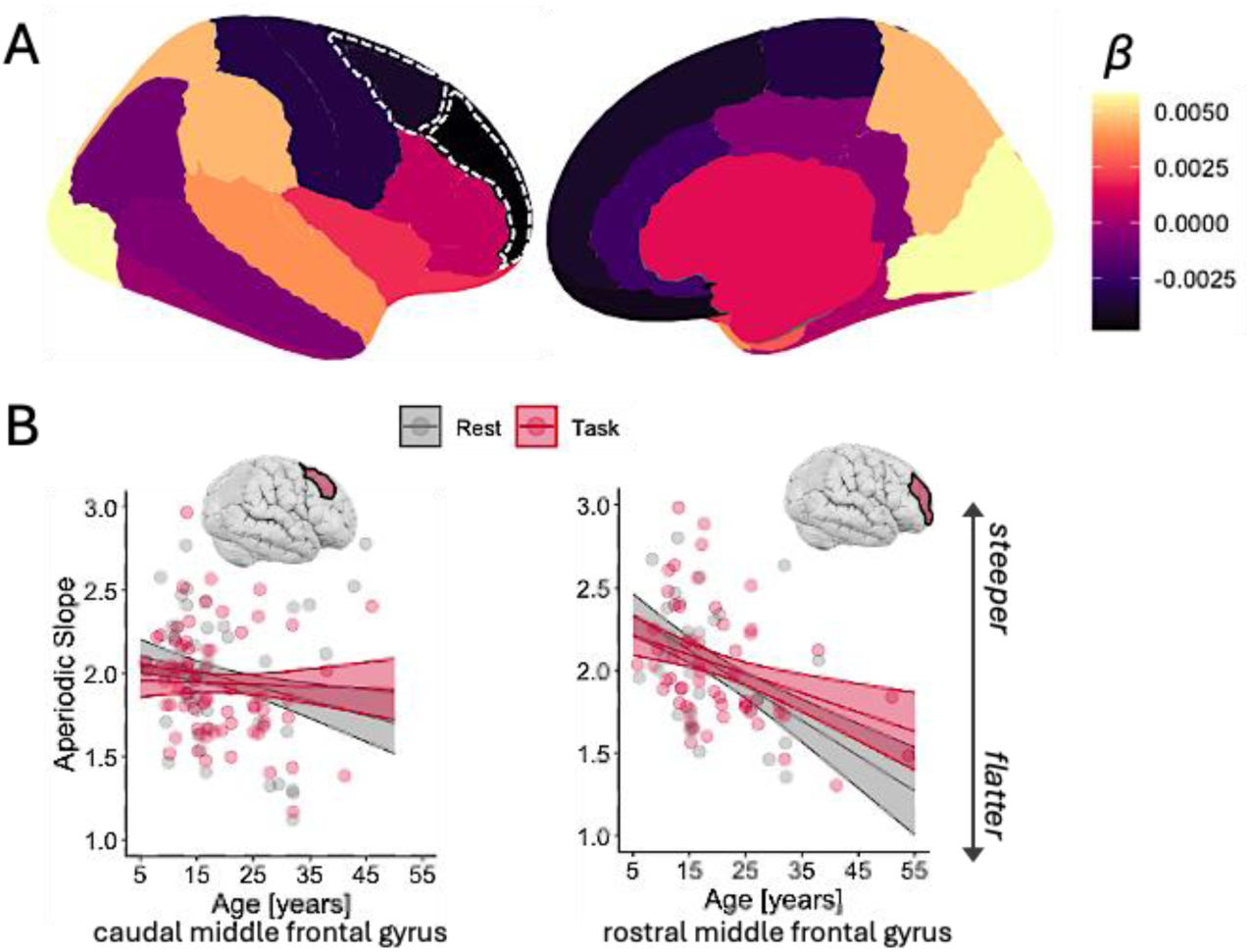
Regions with a significant interaction between age and attentional state on aperiodic activity. **(A)** Brain-wide age and condition interactions on regional aperiodic slopes. Regions with statistically significant interactions between age and attentional state (*FDR* < 0.05) are indicated by dashed borders. **(B)** Scatterplots illustrating interactions between age (x-axis; in years) and attentional state (red = task-based; gray = task-free) on the aperiodic slope (y-axis; higher values denote a steeper slope) in regions with statistically significant interactions. Individual data points represent single participant data averaged across channels for each representative ROI. Shading shows 83% CIs.

### Task-based aperiodic activity in association cortices predicts individual memory outcomes

Having demonstrated that memory performance improves with age, with marked variability among adolescents (Figure 1C), we examined whether age interacts with regionally specific task-based and task-free aperiodic slopes, respectively, to predict memory performance. For each analysis, we fit a general linear model to recognition accuracy and regressed the estimates onto age (in years) and aperiodic slopes (task-based or task-free), and the interaction of age and slope. For task-based slopes (after accounting for the unique contribution of age) we observed an age × slope interaction in rMFG (*β* = 0.02 [95% CI = 0.009, 0.04], SE = 0.01, *p*_adj_ = 0.04; age: *β* = -0.02 [95% CI = -0.06, -0.002], SE = 0.01, *p*_adj_ = 0.14; slope: *β* = -0.76 [95% CI = -1.18, -0.41], SE = 0.23, *p*_adj_ = 0.003; *R*^2^ = .40; Figure 5). In rMFG, memory performance increased with age and an age-related flattening of the aperiodic slope. Although overall steeper slopes were observed in children, relatively flatter slopes in children and adolescents were associated with relatively superior memory. There were no significant main effects of the task-free slope or interactions between the task-free slope and age on memory performance (all *p* >.05). For visualization of model diagnostics, see Figures S6 and S7, and for main effects of task-based and task-free slopes on memory, see Figure S8 and Figure S9, respectively.

**Figure 5.**
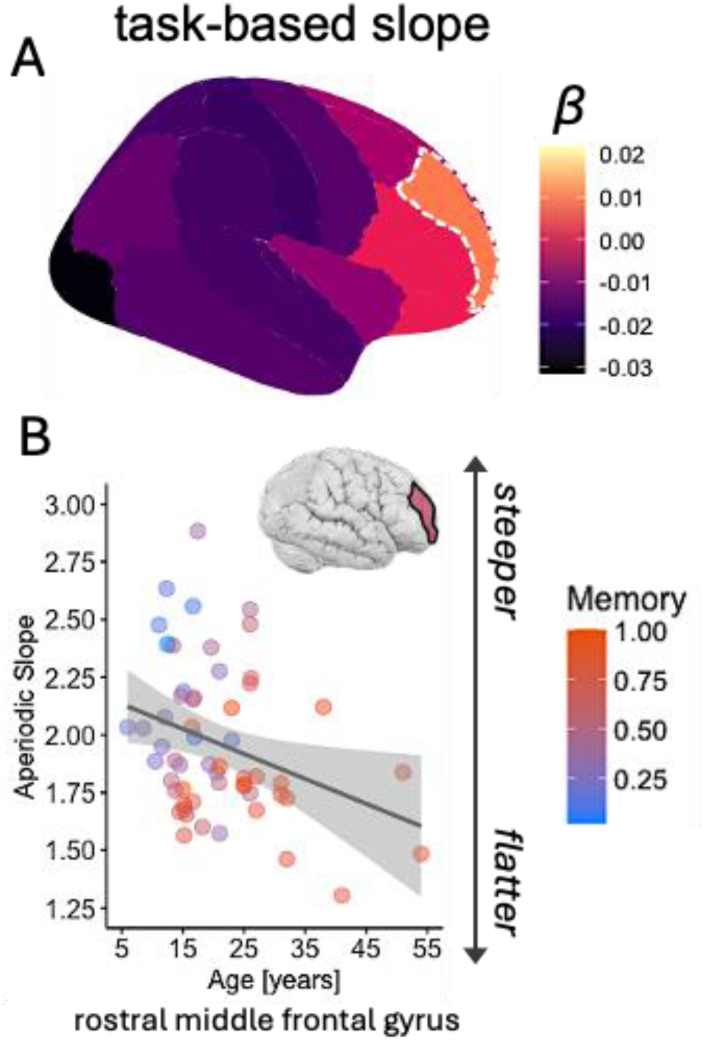
Task-based aperiodic slopes in MFG predict age-related differences in memory performance. **(A)** Brain-wide slope and age interactions on memory, with rMFG demonstrating statistically significant interactions between the task-based slopes and age (*p* < 0.05) on memory performance, with this indicated by dashed borders. **(B)** Scatterplot illustrating interactions between task-based rMFG slopes (y-axis; higher values denote a steeper slope) and age (x-axis; in years) on memory (z-scale; warmer colors denote higher memory recognition accuracy). Individual data points represent single participant data averaged across channels for each representative ROI. Shading shows the standard error.

Taken together, our results elucidate how age-dependent effects on aperiodic slopes in PFC predict individual memory performance. These effects were evident exclusively during task-based states, thus linking aperiodic activity during attention to to-be-remembered visual information to an individual’s memory for that information. From this perspective, the aperiodic slope may serve as a key marker of typical and atypical memory development..

### Gray matter volume and age interact to predict aperiodic activity in select brain regions

Thus far, we have established that aperiodic slopes in PFC differ by age and attentional state and predict age-related variability in memory outcomes, whereas slopes in sensorimotor regions do not differ by attentional state or predict age-related variability in memory outcomes. Last, we focus on structure-function relationships. Before testing hypothesis d, that age-related differences in aperiodic activity are modulated by regional GMV, we sought to replicate previous reports of age-related reductions in regional GMV (Bethlehem et al., 2022; Gogtay et al., 2004; Groeschel et al., 2010). Having demonstrated that global GMV decreases with age in our cohort (Figure 1C), we examined GMV by ROI. We implemented a linear mixed-effects model, regressing region and age onto GMV, treating participants as random intercepts. Our model confirmed a significant main effect of region (χ2(19) = 11081.98, *p* ≤ 0.001) and revealed an age × region interaction (χ2(19) = 92.05, *p* ≤ 0.001; Figure 5). Gray matter volume was reduced with age in lateral orbitofrontal cortex (*r* = -0.61, *p* < 0.001, 95% CI = [-0.77, -0.35]), mOFC (*r* = -0.60, *p* = 0.001, 95% CI = [-0.81, -0.27]), rMFG (*r* = -0.38, *p* = 0.007, 95% CI = [-0.61, -0.11]), cMFG (*r* = -0.42, *p* <.001, 95% CI = [-0.60, -0.20]), inferior frontal gyrus (*r* = -0.35, *p* = 0.01, 95% CI = [-0.57, -0.09]), superior temporal cortex (*r* = - 0.45, *p* < 0.001, 95% CI = [-0.64, -0.22]), middle temporal cortex (*r* = -0.31, *p* = 0.01, 95% CI = [- 0.51, -0.07]), posterior cingulate cortex (*r* = -0.36, *p* = 0.04, 95% CI = [-0.62, -0.02]), and inferior parietal cortex (*r* = -0.54, *p* < 0.001, 95% CI = [-0.70, -0.33]; Figure 6A and 6B). These results replicate previous reports of age-related reductions in GMV in association cortices starting in childhood (Bethlehem et al., 2022; Gogtay et al., 2004; Groeschel et al., 2010).

**Figure 6.**
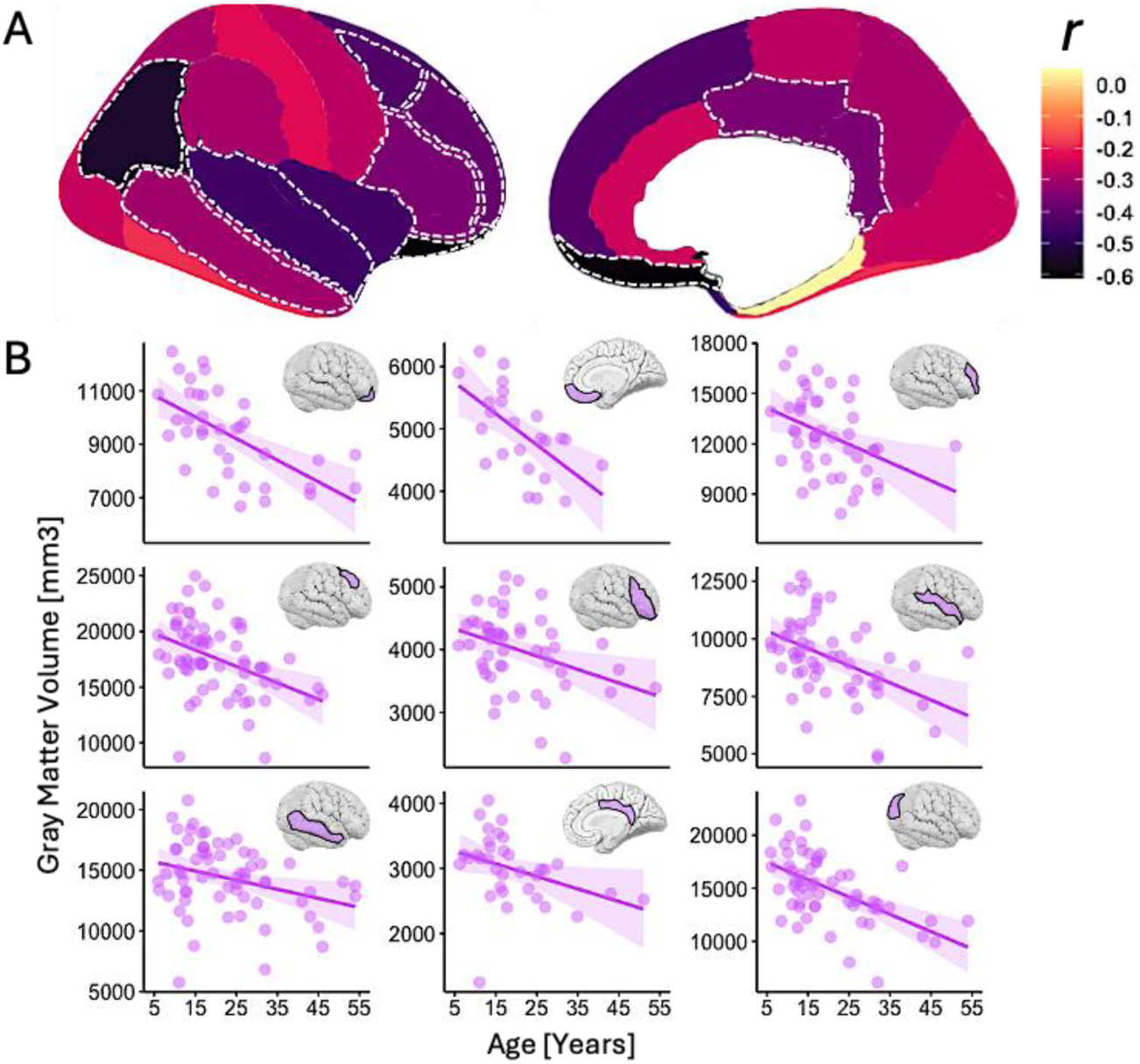
Regions with a significant effect of age on GMV. **(A)** Brain-wide correlations (Pearson *r*) between regional GMV (mm^3^) and age (in years). Warmer colors/higher values indicate positive correlations and cooler colors/lower values indicate negative correlations. Note that the area corresponding to subcortical space is white as no analysis of subcortical GMV was performed. Regions with statistically significant correlations (*p* < 0.05) are indicated by dashed borders. **(B)** Scatterplots illustrating relationships between GMV (y-axis) and age (x-axis) in regions with statistically significant correlations.

To test whether age-related differences in aperiodic activity are modulated by regional GMV, we fit mixed-effects models to task-based aperiodic activity and regressed these estimates onto age (in years), GMV, and the interaction between age and GMV. All models were fit with by-participant and by-task random intercepts, with channel nested under participant. We revealed age × GMV interactions on task-based aperiodic slopes in posterior cingulate cortex (*β* = -4.83 × 10^-5^ [95% CI = 7.58 ×10^-5^, 2.05 × 10^-5^], SE = 1.43 × 10^-5^, *p*_adj_ = 0.04; age: *β* = 0.12 [95% CI = 0.05, 0.20], SE = 0.04, *p*_adj_ = 0.04; GMV: *β* = 0.0006 [95% CI = 0.0002, 0.001], SE = 0.0002, *p*_adj_ = 0.16) and postcentral gyrus (*β* = -1.08 × 10^-5^ [95% CI = -1.79 × 10^-5^, -3.80 × 10^-6^], SE = 3.57 × 10^-6^, *p*_adj_ = 0.04; age: *β* = 0.10 [95% CI = 0.03, 0.17], SE = 0.03, *p*_adj_ = 0.04; GMV: *β* = 0.0001 [95% CI = 1.81 × 10^-5^, 2.51× 10^-5^], SE = 5.93 × 10^-5^, *p*_adj_ = 0.28; Figure 7). In posterior cingulate cortex and postcentral gyrus, while there was no relationship between task-based slopes and GMV in children, flatter slopes were associated with higher GMV in adults (for visualizations of the main effects of age and GMV on aperiodic slopes, see Figures S10 and S13, respectively; for model diagnostics, see Figures S11 and S12).

**Figure 7.**
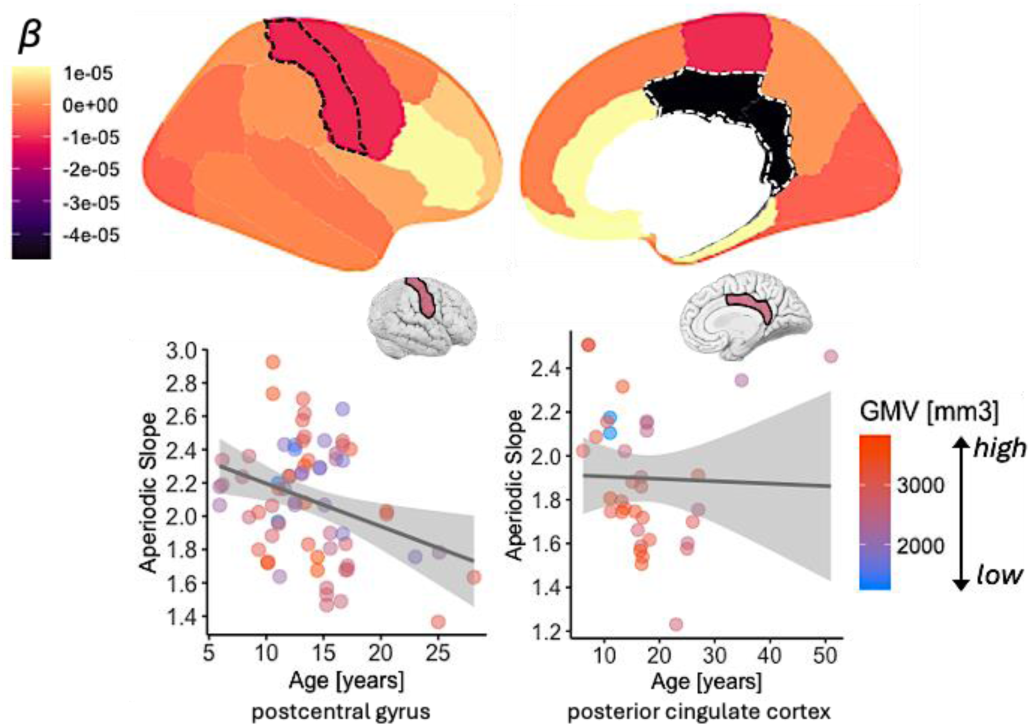
Regions with a significant interaction between age and GMV on the aperiodic slope. **(A)** Top row: Brain-wide GMV and age interactions on regional aperiodic slopes. Regions with statistically significant interactions between age and GMV (*FDR* < 0.05) are indicated by dashed borders. Bottom row: scatterplots illustrating interactions between age (x-axis; in years) and GMV (z-scale; warmer colors denote higher GMV) on the aperiodic slopes (y-axis; higher values denote a steeper slope) in regions with statistically significant interactions. Individual data points represent single participant data averaged across channels for each representative ROI. Shading shows the standard error.

## Discussion

We mapped aperiodic activity – a marker of neural noise – from childhood to late middle adulthood. Our findings demonstrate: (I) a gradient of slopes from inferior lateral to superior medial regions, suggesting reduced neural noise in inferior lateral temporal regions and increased neural noise in superior medial frontal regions (Figure 2); (II) a U-shaped relationship in slopes by age, suggesting increased neural noise into young adulthood followed by reduced neural noise into middle adulthood (Figure 3); (III) a flattening of PFC slopes with advancing age, with more pronounced flattening in task-free states, suggesting that age-related increases in neural noise are task-dependent (Figure 4); (IV) PFC-derived aperiodic slopes during task-based states predict age-related variability in memory (Figure 5); and (V) lower GMV is associated with steeper slopes across age in association and sensorimotor cortices (Figure 7). In sum, these findings reveal regional and attentional differences in neural noise from early childhood to late middle adulthood and establish the balance of neural noise in PFC as a mechanism of memory development (for a schematic summary of the main results, see Figure 8).

**Figure 8.**
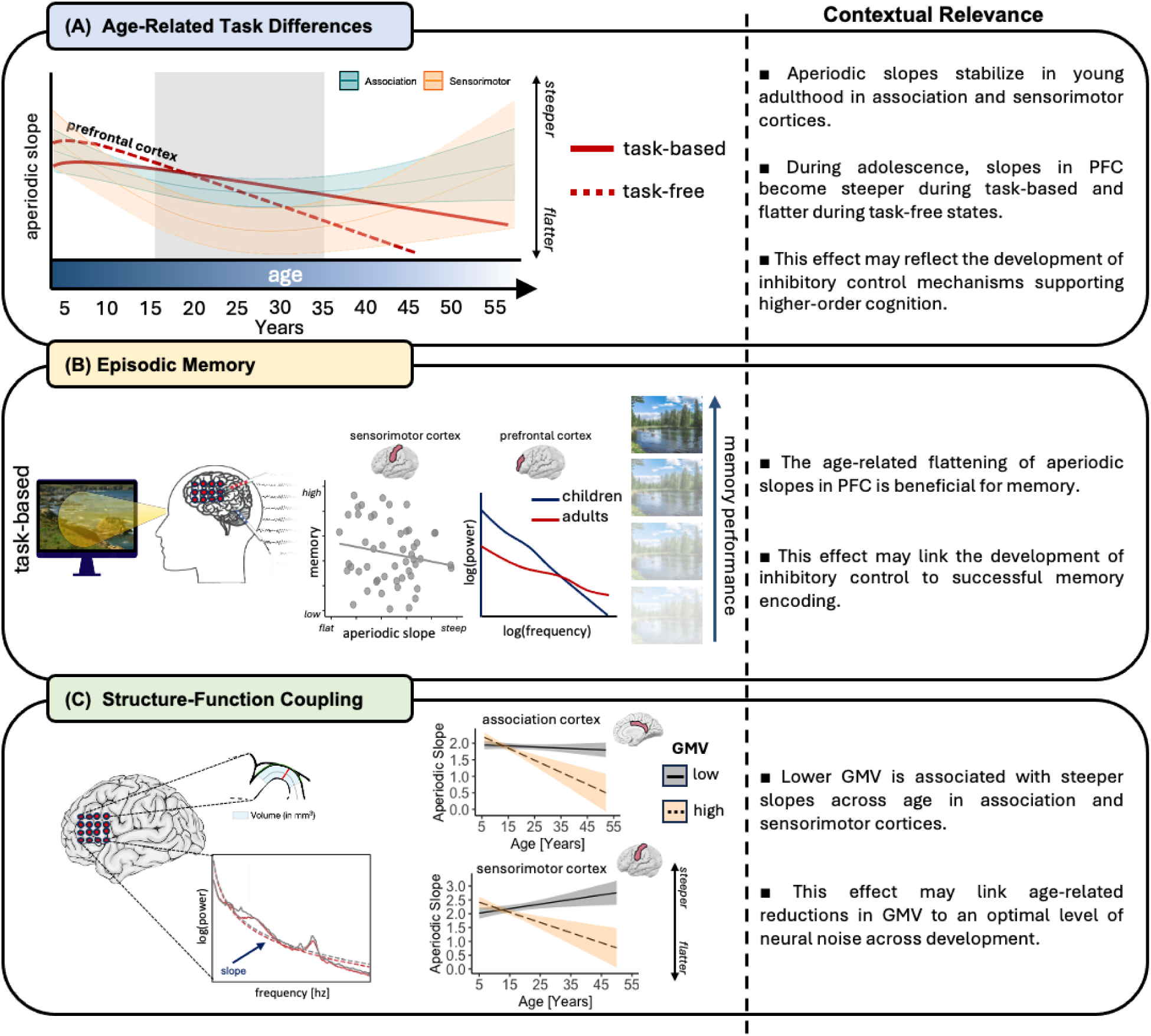
Aperiodic activity stabilizes in young adulthood, differs by age and attentional state, predicts age-related variability in episodic memory, and is associated with age-related variability in GMV. **(A)** Aperiodic slopes in sensorimotor (orange) and association (teal) cortices flatten from age 5 – 25 years and steepen thereafter. Note that the flattening is more pronounced in sensorimotor than association cortices in adolescence and young adulthood (gray shading). Regarding attentional state (i.e., task-based vs. task-free) differences in aperiodic activity, in PFC, task-free (dashed red) slopes are steeper (i.e., less neural noise) than task-based (solid red) slopes in children, and the inverse is observed in adults. Effects reverse at approximately ∼18 – 20 years of age, likely reflecting the development of control. **(B)** PFC-derived aperiodic slopes during task-based but not task-free states predicted age-related variability in memory performance, whereby the age-related flattening of aperiodic slopes was associated with age-related improvements in memory. Flatter sensorimotor cortical slopes were not associated with better memory performance after accounting for age. **(C)** Modeling the relationship between brain volume and aperiodic slopes revealed similar age-related differences in structure-function coupling. In both posterior cingulate cortex and postcentral gyrus, slopes were steeper in childhood regardless of GMV; in adolescence and adulthood, lower GMV was associated with steeper slopes and higher GMV was associated with flatter slopes.

### Aperiodic activity stabilizes in association and sensorimotor cortices in adulthood

The spatiotemporal patterning of cortical maturation progresses from sensorimotor to higher-order association cortices, characterized by heightened plasticity in late-maturing association regions, potentially influencing higher-order cognition in adulthood (Sydnor et al., 2021). Based on these observations, we hypothesized that aperiodic activity would follow similar developmental trajectories, such that it would stabilize during adolescence in sensorimotor cortices and during young adulthood in association cortices. Indeed, recent work (Momi et al., 2025) emphasizes that, while sensorimotor regions exhibit a more localized and intrinsic activation pattern, indicative of a more segregated and stable E/I balance, association regions show more integrated and interconnected activity, with a greater reliance on recurrent feedback loops, reflecting a more dynamic and finely tuned E/I balance that develops later in life. Consistent with our hypothesis, we revealed that the aperiodic slope flattens from childhood to young adulthood in association cortices. However, contrary to dominant models of brain development based on structural measures (Gogtay et al., 2004; Grydeland et al., 2019; Sydnor et al., 2021), we found that aperiodic activity in sensorimotor cortices does not stabilize until young adulthood. We further revealed that the magnitude of flattening is greater in sensorimotor than association cortices during adolescence and young adulthood, pointing to developmental dissociation. Our findings establish that the development of aperiodic activity in sensorimotor regions does not mirror the development of cortical structure and suggest that developmental differences in neural noise in sensorimotor regions follows a protracted trajectory into adulthood.

### Attention modulates aperiodic activity by age in prefrontal cortex

Scalp-EEG studies have consistently demonstrated an age-related flattening of the aperiodic slope, often with a frontal-central distribution (Bornkessel-Schlesewsky, Alday, et al., 2022; Favaro et al., 2023; McSweeney et al., 2023; Merkin et al., 2023; Ouyang et al., 2020; Schaworonkow & Voytek, 2021; Thuwal et al., 2021). To our knowledge, only one iEEG study has examined age-related aperiodic slope variability, demonstrating an age-related flattening of task-based slopes in the visual cortex of 15 individuals aged 15 – 53 years (Voytek et al., 2015). Little is known regarding regional differences in the slope. We found that subregions of PFC, namely caudal and rostral MFG, exhibit a flattening of the aperiodic slope across age. We further reveal that the age-related flattening of the slope is modulated by attentional state, with less pronounced flattening for task-based relative to task-free states. This finding can be interpreted in the context of PFC control: a central role of the PFC is to exert cognitive control in the service of behavior, partially by modulating activity in regions further upstream, such as visual cortex and MTL (Gazit et al., 2020; E. K. Miller & Cohen, 2001; Noudoost & Moore, 2011). The difference between task-states also emerges at roughly 18 to 20 years of age, revealing the aperiodic slope as a potential marker of the development of cognitive control in adolescence, and mirroring the development of domain-general cognitive control (Tervo-Clemmens et al., 2023). Functionally, steeper on-task slopes, suggesting reduced neural noise, have been proposed to reflect the maintenance of top-down predictions (Cross et al., 2022; Dave et al., 2018) and support information integration (Bornkessel-Schlesewsky, Sharrad, et al., 2022; Sheehan et al., 2018). By contrast, flatter slopes have been associated with slower processing speed (Ouyang et al., 2020), and poorer visual working (Donoghue et al., 2020) and visuomotor (Immink, Cross et al., 2021) memory, albeit these studies analyzed task-free slopes. Our findings suggest that the PFC gains flexibility in control with age, exerting increased control during the processing of external, task-relevant information.

### Aperiodic activity during memory encoding predicts subsequent memory performance

Do age-related differences in aperiodic activity predict age-related differences in memory? Prior work on aperiodic activity has reported mixed findings in relating the slope and offset to various aspects of cognition. Steeper task-free slopes have been associated with faster reaction times in young adults and improved recognition accuracy during initial learning (Immink, Cross et al., 2021). However, in the same study, flatter slopes and higher offsets were associated with improved recognition with increasing task exposure. In a similar study with young adults, flatter task-free slopes and higher offsets were associated with improved decision-making performance (Dziego et al., 2023). Of the studies examining task-based aperiodic activity, flatter slopes have been associated with improved learning of an artificial language in young adults aged 18 – 40 years (Cross et al., 2022), but lower working memory performance with age from 15 – 53 years (Voytek et al., 2015). Critically, past work has either focused on task-based or task-free aperiodic activity and cognition without accounting for differences between task-states, and it is unknown how task-based differences in localized brain regions relate to behavior by age.

Here, we overcame this limitation by mapping task-based and task-free aperiodic slopes by age to behavior on a region-by-region basis. We revealed an interaction between aperiodic slopes and age on memory in MFG. In MFG – a region core to executive functions and cognitive control and which undergoes protracted development (Fuster, 2002; Ridderinkhof et al., 2004) – children with steeper slopes exhibited worse memory performance. This finding suggests that too little noise (Donoghue et al., 2020; Voytek et al., 2015) in MFG during childhood may hinder attentional control. Indeed, ADHD-diagnosed, medication naïve children exhibit steeper slopes than their typically developing counterparts (Robertson et al., 2019), as do individuals with schizophrenia (Molina et al., 2020; Peterson et al., 2023), suggesting that even greater reductions in neural noise in childhood results in inefficient neural communication and disrupted coordination, manifesting in poorer memory outcomes.

As individuals age, structural and functional changes in MFG (i.e., synaptic pruning, changes in neurotransmitter levels [GABAergic interneurons, glutamate]; (Kolk & Rakic, 2022), likely lead to a flattening of aperiodic slopes (McKeon et al., 2024; Sukenik et al., 2021). Flatter slopes have been likened to increased neural noise (Bornkessel-Schlesewsky, Alday, et al., 2022; Bornkessel-Schlesewsky, Sharrad, et al., 2022; Dave et al., 2018; Voytek et al., 2015), due to increased levels of aberrant neural firing in the absence of a slower modulatory oscillation (Voytek et al., 2015; Voytek & Knight, 2015). We observed that flatter slopes in MFG during adulthood were less related to memory outcomes than in children, likely due to the emergence of compensatory neural recruitment and altered cognitive strategies (Braver et al., 2009; Cabeza et al., 2018; Spreng & Turner, 2019).

Interestingly, we did not observe significant relationships between task-free slopes and memory performance. This apparent discrepancy with past findings can likely be explained by differences in experimental task designs and inter-regional source mixing inherent to scalp-EEG, where signals from multiple cortical areas are mixed due to volume conduction (Musall et al., 2014; Palva et al., 2018). Scalp-EEG, with its relatively low spatial resolution, could mask region-specific relationships between aperiodic slopes and behavior, explaining discrepancies with previous findings. Although source localization techniques can help mitigate these issues, they are limited in resolving precise cortical sources (Buzsaki, 2006; Nunez & Srinivasan, 2006). Further, previous work has focused on mapping intrinsic, task-free aperiodic activity onto trait-like measures of cognition (e.g., processing speed, verbal ability; (Euler et al., 2024; Montemurro et al., 2024; Pi et al., 2024) or tasks that do not measure episodic or working memory (Bornkessel-Schlesewsky, Sharrad, et al., 2022; Dziego et al., 2023; Immink et al., 2021). Our findings demonstrate that aperiodic activity during the encoding of visual stimuli predicts recognition of those stimuli, a direct relationship that did not survive on a region-by-region basis with intrinsic (i.e., task-free) activity.

### Aperiodic activity is associated with age-related variability in gray matter volume

Finally, having established that aperiodic activity differs by age and attentional state and that task-based aperiodic activity predicts memory outcomes, we mapped task-based aperiodic activity onto GMV across age. We reveal that in posterior cingulate cortex (PCC), aperiodic slopes are not dependent on GMV during childhood. However, as individuals age, the relationship between GMV and aperiodic slopes becomes pronounced. Specifically, individuals who maintain higher GMV in the PCC exhibit significant flattening of aperiodic slopes over time. In contrast, those with lower GMV in the PCC during development show stable, steep aperiodic slopes, meaning that their slopes do not differ with age and remain relatively steep across development.

Mechanistically, this pattern may reflect differential cortical maturation and the influence of GMV on neural network stability. In individuals with higher GMV, the gradual stabilization of synaptic networks may facilitate a more balanced cortical state, resulting in flatter aperiodic slopes as they age. In those with lower GMV, however, the lack of such stability may prevent the flattening of the aperiodic slope, leading to a persistently steeper slope despite developmental differences. These findings suggest that early cortical changes in GMV, particularly in the PCC, may be key to understanding the long-term stability of aperiodic neural dynamics and their relationship to cognitive processes such as memory. This should be further elucidated in future work.

Similarly, in postcentral gyrus (i.e., primary motor cortex), we observed that lower GMV is associated with flatter slopes during childhood and steeper slopes during adulthood. This is in addition to the finding that sensorimotor cortical development – as indexed by neural noise – stabilizes during young adulthood, challenging models of early sensorimotor development based on cortical structure (Bethlehem et al., 2022; Larsen et al., 2022). Future research should further examine relationships between aperiodic activity and brain structure to elucidate the mechanisms by which structure-function development impacts the development of higher-order cognition.

### Limitations and future directions

We have revealed regional age-related variations in aperiodic neural activity dependent upon task-state. Our findings suggest that brain development may be best understood as a diverse set of regionally independent trajectories, partially indexed by aperiodic activity. However, as iEEG data are cross-sectional, we were unable to follow these putative trajectories through time. A critical next step will be to establish the potential utility of aperiodic activity in elucidating longitudinal changes in regional structure-function relationships (Ofen et al., 2019). As such, future studies, focusing *a priori* on the regions we identified (e.g., MFG), could capitalize on the spatio-temporal precision and capacity to perform multi-visit longitudinal studies with, for example, MEG.

While our cohort is representative of typical development and the use of iEEG affords precise spatiotemporal precision, iEEG samples are comprised of pharmacoresistant epilepsy patients, potentially limiting the generalizability of our findings (Johnson & Knight, 2023). For this reason, it is important to note that our sample demonstrated typical age-related gains in memory performance and age-related differences in global GMV (Figure 1C), both of which are consistent with healthy cohorts (Bethlehem et al., 2022). An additional limitation is the relatively lower representation of older individuals within our sample, a common observation in iEEG investigations, and the relatively lower representation of patients with task-free (*n* = 65) compared to task-based (*n* = 81) data. We also acknowledge that there was low sampling of certain brain regions (see Tables S2 and S3 for channel count by region), which may reduce the generalizability of some results. Low sampling reduces statistical power, potentially masking meaningful neural activity or introducing biases in region-specific analyses (Johnson, Yin, et al., 2022; Johnson & Knight, 2023). To address this limitation, we caution readers against over-interpreting the results for regions the amygdala, insula, and superior parietal cortex. Further, in our analysis, we focused on the frequency range of 1–60 Hz and did not specifically account for the potential influence of a ’knee’ in the PSD, typically observed around 75 Hz (K. J. Miller et al., 2009). This knee can result in a shift in the PSD’s scaling behavior, which may affect the aperiodic slope values if not addressed in the fitting process. Future research should prioritize obtaining more balanced sampling across these regions, possibly by using even larger, more diverse datasets, and in using algorithms that account for potential “knees” in the PSD (Donoghue et al., 2020). Nonetheless, the current results underscore maturation within MFG, and this effect was present across our entire age range of ∼5 to 54 years. To obtain larger samples across age, future research may seek to increase sample sizes through multi-site collaboration and data sharing (Johnson, Yin, et al., 2022; Johnson & Knight, 2023).

We also found no significant age-related difference in aperiodic activity in the hippocampus in relation to attentional state, or in predicting individual memory performance. Given that our study examined memory, these results may be somewhat surprising. However, it is possible that the development of oscillatory (i.e., periodic) activity in the hippocampus exhibits effects related to attentional state and memory outcomes, consistent with ample literature on hippocampal theta oscillations (Herweg et al., 2020; Lega et al., 2012; Yin et al., 2023). Future research should directly investigate this hypothesis. Lastly, with our task-based versus task-free contrast as a starting point, future research may also aim to examine additional attentional states, such as sleep versus wake states. The aperiodic slope and offset systematically shift as a function of sleep stage, which has recently been shown to differ across development (Favaro et al., 2023). However, it is unknown whether there are region-specific differences in sleep-based aperiodic activity, whether these regional differences relate to the development of higher-order cognition, and whether sleep-based aperiodic activity changes concomitantly with wake-related aperiodic dynamics.

### Implications

Historically, neuroscientific research has predominantly focused on young adults aged 18-40 years, largely overlooking the influence of age on brain dynamics. This practice has resulted in a significant knowledge gap regarding brain development. Addressing this gap is crucial due to its profound clinical implications across various domains, including neurodevelopmental disorders, traumatic brain injury, stroke, age-related cognitive decline, and neurodegenerative diseases, as well as advancements in neural prosthetics for injury, stroke, or disease management. Our study addresses this knowledge gap by elucidating the trajectory of aperiodic electrophysiological dynamics and their associations with brain structure and memory across development, from childhood into late middle adulthood. Previous attempts to characterize these dynamics have been constrained by limitations in imprecise spatiotemporal measurements and relatively small sample sizes. To overcome these challenges, we adopted a comprehensive approach. Firstly, we employed iEEG to delineate developmental neurophysiology with exceptional precision. Secondly, we applied sophisticated analyses of aperiodic components in iEEG data to establish novel connections between aperiodic activity and developmental variations in memory. Thirdly, we explored the relationship between aperiodic components and GMV. Lastly, we leveraged an exceptionally large iEEG dataset to detect subtle effects that may have been undetected in smaller cohorts.

Understanding how cortical maturation influences memory encoding processes is also fundamental to cognitive function and daily performance, given well-documented changes in brain structure and function over the lifespan. Furthermore, elucidating the impact of brain development on memory formation across different life stages holds promise for early detection and intervention strategies targeting the emergence of both neurodevelopmental disorders and age-related memory decline. Identifying markers of healthy brain development and aging is crucial for detecting dysfunction in age-related pathologies, which often manifest gradually over many years before exhibiting overt behavioral symptoms. In this context, our findings may contribute to the prevention or delay of pathological aging, offering significant health benefits, particularly considering the limitations and risks associated with current pharmacological treatments. Additionally, our study lays the groundwork for investigating memory dysfunction in psychiatric disorders, many of which emerge during adolescence and young adulthood, and which show deviations in aperiodic activity from healthy populations (Earl et al., 2024; Fernandez & Garner, 2007; Pani et al., 2022; Shuffrey et al., 2022).

### Conclusions

We reveal that aperiodic neural activity follows the same developmental time course across young adulthood in both sensorimotor and association cortices, challenging models of early sensorimotor development based on measures of brain structure. We also isolate attentional state and age-related differences in the aperiodic slope to PFC, demonstrating that task-based slopes are steeper, reflecting less neural noise, and that this difference emerges during adolescence. We further establish the functional role of PFC-derived slopes in memory, revealing that age-related improvements in memory outcomes are associated with the age-related flattening of aperiodic slopes. Lastly, we characterized, for the first time, the relationship between age-related differences in aperiodic activity and brain structure, identifying region-specific trajectories in structure-function relationships during development. Taken together, our findings establish brain-wide maps in aperiodic neural activity, its relation to age-related variability in memory, and novel structure-function relationships, findings which are critical for understanding brain development and aging in both health and disease.

## Methods

### Participants

Participants were 101 neurosurgical patients aged 5.93 – 54.00 years (63 males; mean age = 19.25) undergoing iEEG monitoring as part of clinical seizure management. Those with major lesions, prior surgical resections, noted developmental delays, or neuropsychological memory test scores <80 were considered ineligible. Patients were recruited from Northwestern Memorial Hospital, the Ann & Robert H. Lurie Children’s Hospital of Chicago, St. Louis Children’s Hospital, University of California, Irvine Medical Center, University of California, Davis Medical Center, University of California, San Francisco Benioff Children’s Hospital, Children’s Hospital of Michigan, and Nationwide Children’s Hospital, University of California, San Diego, Rady Children’s Hospital, Mount Sinai Hospital, California Pacific Medical Centre, and University of California, San Francisco Medical Centre. Written informed consent was obtained from participants aged 18 years and older and from the guardians of participants aged under 18 years. Written assent was obtained from participants aged 13 – 17 years and oral assent was obtained from younger children. All procedures were approved by the Institutional Review Board at each hospital in accordance with the Declaration of Helsinki. Given that electrode positioning in these participants was based on clinical necessity rather than for experimental reasons, *a priori* power analyses were not performed. Human iEEG research is limited by the availability of neurosurgical patients. From this perspective, the majority of iEEG work has been based on relatively small sample sizes and could not consider age-related or other sources of inter-individual variability (Johnson & Knight, 2023).

### Experimental design

Task-based iEEG data were derived from the encoding phase of two visual memory recognition tasks that have been used extensively to study memory in adults and children across neuroimaging modalities, including iEEG. In the blocked-trial paradigm, participants encode a set of 40 indoor and outdoor scenes and classify each as indoor/outdoor in preparation for a self-paced old/new recognition test of all 40 studied scenes intermixed with 20 new scenes as foils (Chai et al., 2010, 2014; Johnson, Tang, et al., 2018; Johnson, Yin, et al., 2022; Ofen et al., 2007, 2012, 2019; Tang et al., 2018; Yin et al., 2020, 2023). In the single trial paradigm, participants encode three shapes in a specific spatiotemporal sequence in preparation for a self-paced old/new recognition test of sequences that match exactly or mismatch on one dimension (i.e., shape identity, spatial position, or temporal order; cf. (Davoudi et al., 2021; Dezfouli et al., 2021; Johnson, Adams, et al., 2018; Johnson, Chang, et al., 2022; Johnson et al., 2017, 2019). Both paradigms use visual stimuli to avoid potential confounds on memory with verbal material in children. The encoding phases of the two paradigms are similar because, in both paradigms, participants encode visual stimuli (3000ms, 500-1500ms inter-trial interval) in preparation for a self-paced, two-alternative forced choice recognition test. We ensured that on-task data reflected task engagement by only analyzing iEEG data during the viewing of stimuli that were attended during encoding, as indexed by a correct indoor/outdoor classification of each scene in the blocked-trial paradigm and correct old/new classification of each sequence in the single-trial paradigm (Johnson, Tang, et al., 2018; Johnson, Yin, et al., 2022; Yin et al., 2020). For a schematic of both visual memory tasks, see Figure S14. For task-free data, participants were instructed to sit quietly with their eyes open, fixating on the center of a computer monitor for five minutes. If no formal task-free task was administered, five minutes of task-free data were taken from natural rest in continuous 24/7 iEEG recordings. This occurred for 27 participants.

### Behavioral analysis

Both visual memory tasks test memory in a two-alternative forced choice design, permitting the use of similar measures of memory performance across tasks. For both tasks, for all participants, we calculated the hit rate (i.e., number of previously studied stimuli that were correctly recognized as old/match out of all studied stimuli) and false alarm rate (number of new stimuli presented that were incorrectly identified as old/match out of all new/mismatched stimuli). Performance accuracy was calculated as hit rate minus false alarm rate to equate measures across memory tasks and correct for differences in an individual’s tendency to respond old/match or new/mismatch, respectively. For a summary of behavioral performance, see Figure 1C.

### iEEG acquisition and pre-processing

iEEG data were recorded at a sampling rate of 200-5000 Hz using Nihon Kohden JE120 Neurofax or Natus Quantum LTM recording systems, which at three sites were interfaced with the BCI2000 software. Data acquired >1000 Hz were resampled to 1000 Hz after the fact. As described below, spectral analysis was performed up to 60 Hz. Thus, the lowest sampling rate of 200 is well over the minimum Nyquist frequency required for analysis (i.e., 2 cycles/frequency = 120 Hz). For consistency, all data from both visual memory tasks and from task-free recordings were pre-processed using the same procedures. Raw electrophysiological data were filtered with 0.1-Hz high-pass and 300-Hz low-pass finite impulse response filters, and 60-Hz line noise harmonics were removed using a discrete Fourier transform. Task-based continuous data were demeaned and epoched into 3s trials (i.e., 0-3s from scene or study sequence onset). Continuous task-free data were also demeaned and transformed into 3s epochs with 25% overlap. All epoched data were manually inspected blind to electrode locations and experimental task parameters. Electrodes overlying seizure onset zones and electrodes and epochs displaying epileptiform activity or artifactual signal (from poor contact, machine noise, etc.) were excluded (mean proportion of rejected epochs = 16.96%, *SD* = 12.51). We employed a bipolar re-referencing strategy, which has been shown to minimize the impact of impedance and electrode size differences between sEEG and ECoG and thus maximize standardization across these two types of recordings (Mercier et al., 2022). Neighboring electrodes within the same anatomical structure were re-referenced using consistent conventions (ECoG, anterior – posterior; sEEG, deep – surface). For ECoG grids, electrodes were referenced to neighboring electrodes on a row-by-row basis. An electrode was discarded if it did not have an adjacent neighbor, its neighbor was in a different anatomical structure, or both it and its neighbor were in white matter. Bipolar referencing yielded virtual channels that were located midway between the original physical electrodes. Data were then manually re-inspected to reject any trials with residual noise. Pre-processing routines used functions from the FieldTrip toolbox for MATLAB (Oostenveld et al., 2011). All results were based on analysis of non-pathologic, artifact-free channels, ensuring that data represented healthy cortical tissue (Rossini et al., 2017).

### Aperiodic neural activity

The irregular-resampling auto-spectral analysis method (Wen & Liu, 2016) (IRASA) was used to estimate the 1/ƒ power-law exponent. IRASA estimates the aperiodic component of neural time series data by resampling the signal at multiple non-integer factors *h* and their reciprocals 1/*h*. As this resampling procedure systematically shifts narrowband peaks away from their original location along the frequency spectrum, averaging the spectral densities of the resampled series attenuates peak components while preserving the 1/ƒ distribution. The exponent summarizing the slope of aperiodic spectral activity is then calculated by fitting a linear regression in log-log space. Using the YASA toolbox (Vallat & Walker, 2021) v.0.6.3, we fit a power-law function to each epoch within the frequency range of 1 – 60 Hz. For each epoch, channel, and task, the inverse slope of the power-law function was taken as the trial-level estimate of the 1/ƒ exponent.

### iEEG localization

Macro-electrodes were surgically implanted for extra-operative recording based solely on clinical need. The electrodes were subdural electrode grids or strips with 10 mm spacing or stereoelectroencephalography electrodes with 5-10 mm spacing. Anatomical locations were determined by co-registering post-implantation computed tomography coordinates to pre-operative magnetic resonance (MR) images, as implemented in FieldTrip (Stolk et al., 2018), FreeSurfer (Fischl, 2012), iELVis (Groppe et al., 2017) or VERA (Adamek et al., 2022). Electrode locations were then projected into standard MNI space and bipolar channel locations (see preprocessing) were projected at the midpoint between their contributing electrodes. Based on these MNI coordinates, the *R* package *label4MRI* v1.2 (https://github.com/yunshiuan/label4MRI) was used to categorize each channel into its corresponding Brodmann area, which were then grouped according to the DKT atlas (Klein & Tourville, 2012).

### Structural imaging and regional gray matter volume

T1-weighted MRI scans were acquired as part of routine preoperative procedures. Parcellation of cortex into regions of interest (ROI) was performed based on standard procedures implemented within FreeSurfer (Fischl, 2012). Regional GMVs were then estimated based on the DKT atlas (Klein & Tourville, 2012). GMV from each ROI was calculated using FreeSurfer (Fischl, 2012). Volumes were calculated for left and right ROIs and averaged across hemispheres for analysis.

### Statistical analysis

Data were imported into *R* version 4.2.3 (R Core Team, 2020) with the aid of the *tidyverse* package (Wickham et al., 2019) and analyzed using linear and nonlinear mixed-effects models fit by restricted maximum likelihood (REML) using *lme4* (Bates, 2010) and *splines* (R Core Team, 2020). *P*-values for region-specific models were estimated using the summary function from the *lmerTest* package, which is based on Satterthwaite’s degrees of freedom (Kuznetsova et al., 2017), and Type II Wald Tests from the *car* package (Fox & Weisberg, 2018) for examination of whole-brain effects (i.e., models which included all ROIs). Effects were plotted using the package gg*effects* (Lüdecke, 2018) and *ggplot2* (Wickham & Wickham, 2016). Spearman correlations were used to assess structure-function relationships without the effect of age, with coefficients used to plot region-specific relationships between aperiodic activity and GMV across the whole brain. Statistical significance was adjusted using the False Discovery Rate with an alpha threshold of .05. Task was entered as an unordered factor using sum-to-zero contrast coding and age was specified as a continuous predictor. Please see Table 1 below for a summary of the main analyses, including the types of models employed and their fixed and random effects structures.

**Table 1.** Summary of main statistical tests including number of models computed, outcome variables, fixed effects, and random effects.

| Analysis | #Models | Type | Outcome | Fixed Effects | Random Effects | FDR |
| --- | --- | --- | --- | --- | --- | --- |
| Association vs sensorimotor | 1 | Nonlinear mixed effects regression | Slope | 1. Age (continuous; one spline, 2 knots)<br>2. Cortex type (categorical) | 1. Participant (intercept)<br>2. ROI (intercept)<br>3. Channel (nested under participant) | No |
| Attentional state | 21 | Linear mixed effects regression | Slope | 1. Age (continuous)<br>2. Condition (categorical) | 1. Participant (intercept)<br>2. Channel (nested under participant)<br>3. Task (intercept) | Yes |
| Memory (task-based) | 21 | General linear regression | Accuracy | 1. Age (continuous)<br>2. Slope (continuous) | NA | Yes |
| Memory (task-free) | 21 | General linear regression | Accuracy | 1. Age (continuous)<br>2. Slope (continuous) | NA | Yes |
| Gray matter volume | 20 | Linear mixed effects regression | Slope | 1. Age (continuous)<br>2. GMV (continuous) | 1. Participant (intercept)<br>2. Channel (nested under participant)<br>3. Task (intercept) | Yes |

*Note:* #Models refers to the number of models performed for each analysis; outcome = dependent variable; Cortex type has two levels of association and sensorimotor; Condition has two levels of task-free and task-based; False Discovery Rate (FDR) corrections were applied to all Analyses apart from “Association vs sensorimotor”, given that one model was computed.

In our preregistration, we specified that we would apply a spline to age to model potential non-linear effects of age on aperiodic activity for each ROI, as well as a random effect of task-free recording type (eyes open vs eyes closed). However, in doing so, models indicated nonconvergence or singular fit. To reduce model complexity, we modeled age as a linear predictor and removed task-free recording type as a random effect in our analysis of each ROI. For analyses testing hypotheses a and b, where we tested differences in association and sensorimotor cortices, we had sufficient power to model nonlinear differences. Specifically, we used linear mixed-effects models with a single spline with two internal knots on age specified using the *splines* package. The spline terms allow for nonlinear relationships between age and aperiodic slopes by fitting piecewise polynomials to the data, which provides more flexibility than a simple linear term. This approach enables the model to capture potential nonlinear patterns in the data, which is necessary to test our hypotheses that slopes would vary by age in childhood and then stabilize in adolescence or adulthood.

Also note that when contrast coding is explicitly described, the need for post-hoc testing is eliminated (for a detailed discussion of contrast coding in linear mixed-effects regressions, please see (Brehm & Alday, 2022). Further, for modeled effects, an 83% confidence interval (CI) threshold was used given that this approach corresponds to the 5% significance level with non-overlapping estimates (Austin & Hux, 2002; MacGregor-Fors & Payton, 2013). In order to isolate outliers for variables that were specified as outcomes (i.e., aperiodic slopes, memory performance), we used Tukey’s method (Tukey, 1977), which identifies outliers as exceeding ±1.5 × inter-quartile range. The packages *ggseg* (Mowinckel & Vidal-Piñeiro, 2020) and *ggsegDKT* were used to generate cortical plots based on DKT atlas nomenclature. Hypotheses a and b were tested using the following formula (note that in all models, asterisks (*) denote interaction terms, while + denotes additive terms. *B*_0_ denotes the intercept of the model, while *B*_1_, *B*_2_ etc. denote the chronological specification of fixed effect parameters):

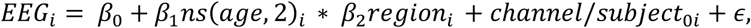

where *EEG* is the aperiodic slope; *age* is age in years modeled with one spline term with two internal knots, and *region* encodes association and sensorimotor cortices; *channel* encodes region-specific channels nested under the random intercept of *participant*, and *participant* is the random intercept term of participant ID. To test hypothesis c, we employed the following model equation on a region-by-region basis:

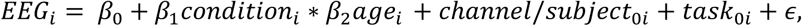

where *EEG* is the aperiodic slope; *condition* encodes task-based and task-free recordings, age is age in years as a linear predictor; *channel* encodes region-specific channels nested under *participant*, and *participant* is participant ID, while *task* is a random intercept encoding whether the recording is derived from the working memory or scene recognition tasks.

Our exploratory analyses focused on relationships between GMV, behavioral performance, and aperiodic slopes derived from task-based and task-free recordings. Here, our primary exploratory research questions were whether:

1. regional age-related variability in aperiodic slopes predicts variability in memory performance, and;
2. regional age-related variability in GMV predicts regional variability in aperiodic slopes.
3. exploratory analyses were examined with general linear models with the following formulae:

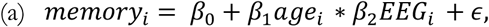

where *memory* is performance on the visual memory task(s), *age* is age in years, and *EEG* is the aperiodic slope from each ROI.

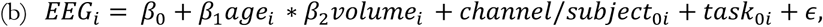

Here, *EEG* is the aperiodic slope; *age* is age in years; *volume* is regional GMV in mm^3^; *channel* encodes ROI-specific channels; *participant* is participant ID; *task* encodes whether the task recording was from the scene recognition or working memory task. As with the other models, each ROI was applied to the model equation described above. *Participant* was modeled as a random effect on the intercept, while *channel* was nested under participant. *Task* were also specified as a random effect on the intercept. Note that in our preregistration, we stated that we would include task (task-based, task-free) in all models examining the interaction between GMV and age on aperiodic activity. However, all models indicated nonconvergence or singular fits. To reduce model complexity, we examined aperiodic activity during task-based states only.

## Acknowledgements

This research was supported in part through the computational resources and staff contributions provided for the Quest high performance computing facility at Northwestern University which is jointly supported by the Office of the Provost, the Office for Research, and Northwestern University Information Technology. We also thank Dr Phillip M. Alday for helpful discussions regarding statistical modeling, Dr. Kurtis I. Auguste for assistance, and the three anonymous reviewers for valuable comments.

