## supplementary material for "The development of aperiodic neural activity in the human brain"

**Table S1.** Summary of association and sensorimotor regions.

| Association | Sensorimotor |
|---|---|
| Caudal Anterior Cingulate | Lateral Occipital Cortex |
| Caudal Middle Frontal Gyrus | Precentral Gyrus |
| Fusiform Gyrus | Postcentral Gyrus |
| Inferior Frontal Gyrus | |
| Inferior Parietal Cortex | |
| Inferior Temporal Cortex | |
| Insula | |
| Lateral Orbitofrontal Cortex | |
| Medial Orbitofrontal Cortex | |
| Middle Temporal Cortex | |
| Parahippocampal Gyrus | |
| Posterior Cingulate Cortex | |
| Rostral Middle Frontal Gyrus | |
| Superior Parietal Cortex | |
| Superior Temporal Cortex | |

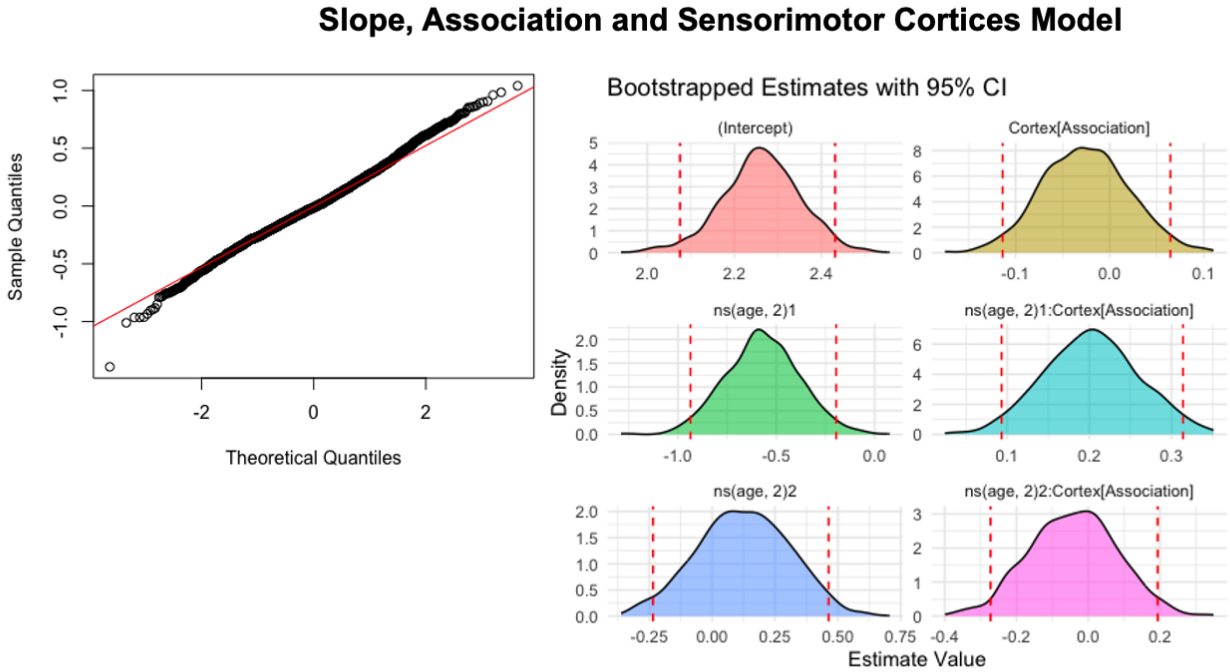


**Figure S1. Residuals and bootstrapped beta coefficients for slope and association vs. sensorimotor model.** Left: Q-Q plot of residuals of the mixed-effects model. Right: Bootstrapped confidence intervals for the intercept, main effects and interactions. Y-axis represents the density of the bootstrapped coefficients and the x-axis represents the estimated beta values. The dashed red lines indicate the 95% confidence interval from the original mixed-model estimates.

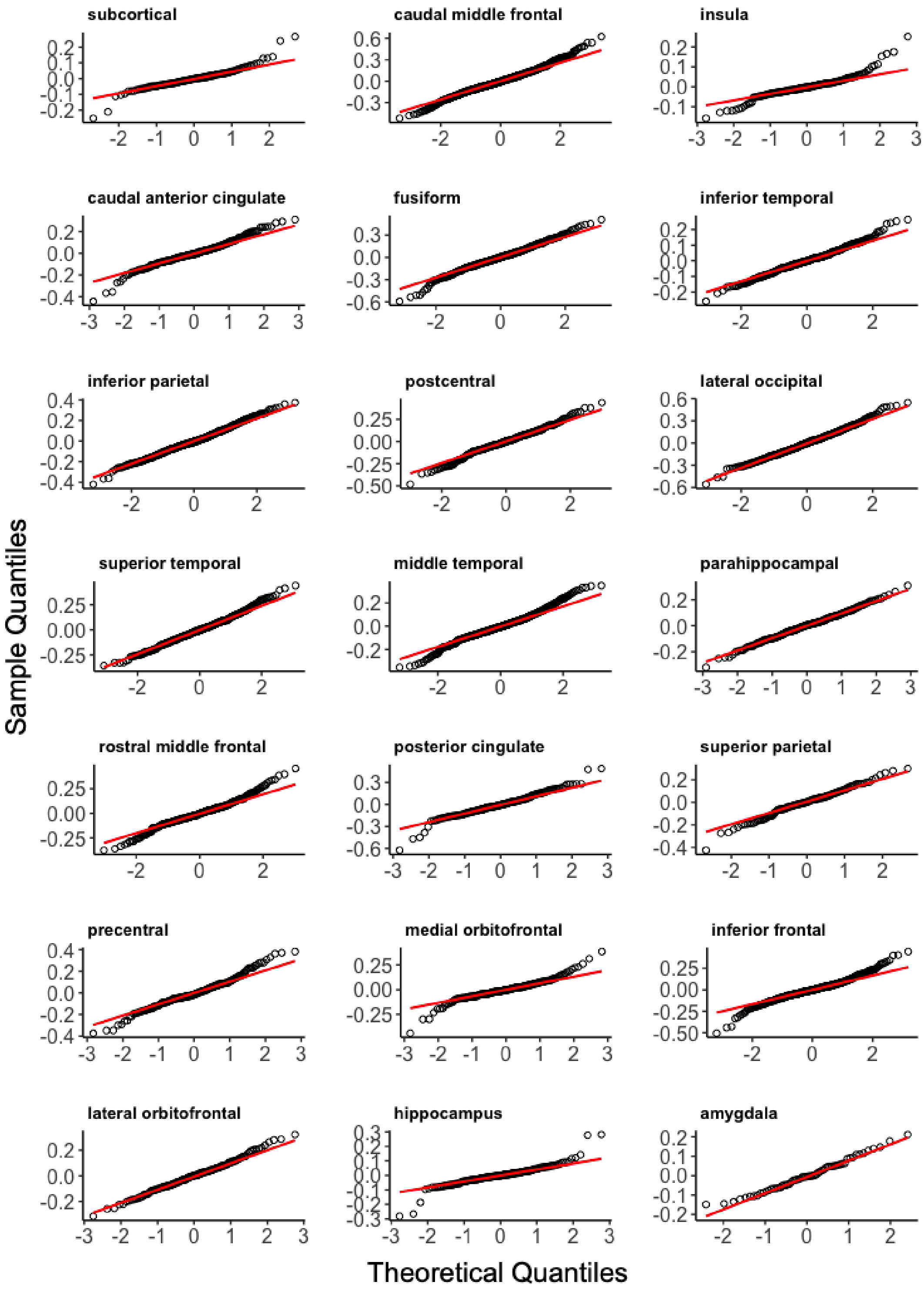


**Figure S2.** Q-Q Plots of residuals for models examining age and attentional state on aperiodic slopes for each ROI.

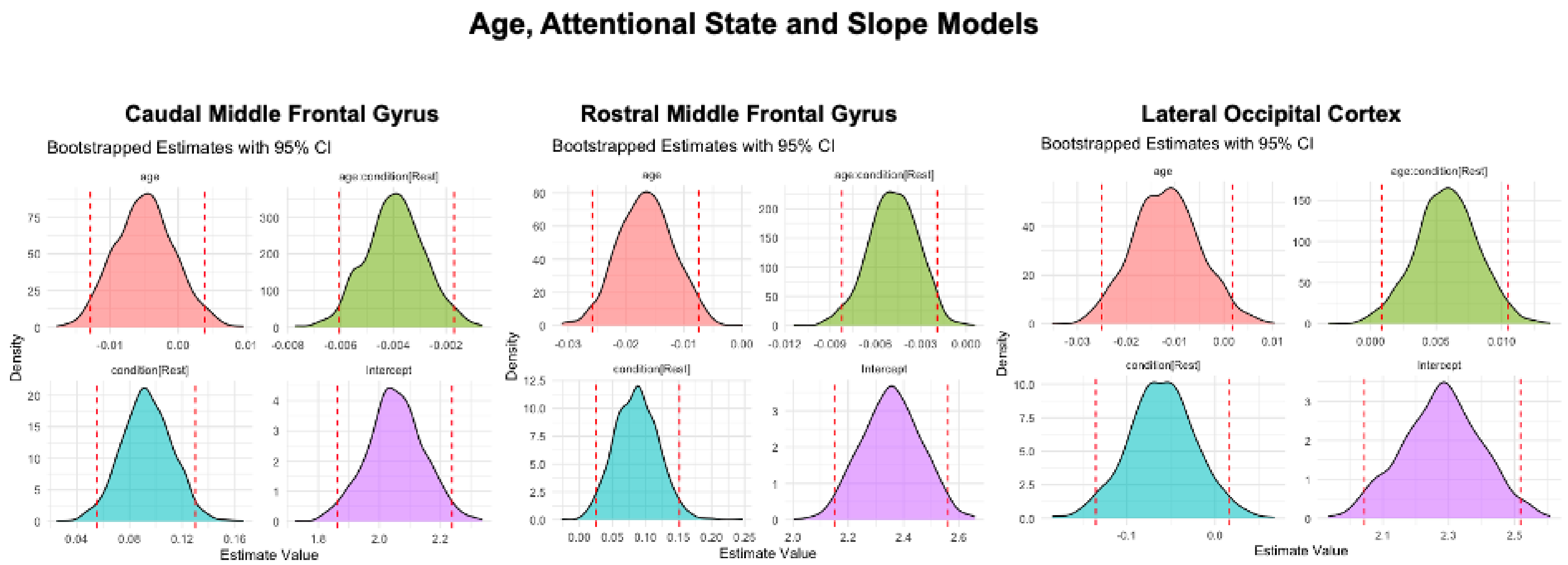


**Figure S3. Bootstrapped beta coefficients for age, attentional state and slope models for the regions that survived multiple comparison correction (left, caudal middle frontal gyrus; middle, rostral middle frontal gyrus; right, lateral occipital cortex).** Bootstrapped confidence intervals for the intercept, main effects, and interactions. Y-axis represents the density of the bootstrapped coefficients and the x-axis represents the estimated beta values. The dashed red lines indicate the 95% confidence interval from the original mixed-model estimates.

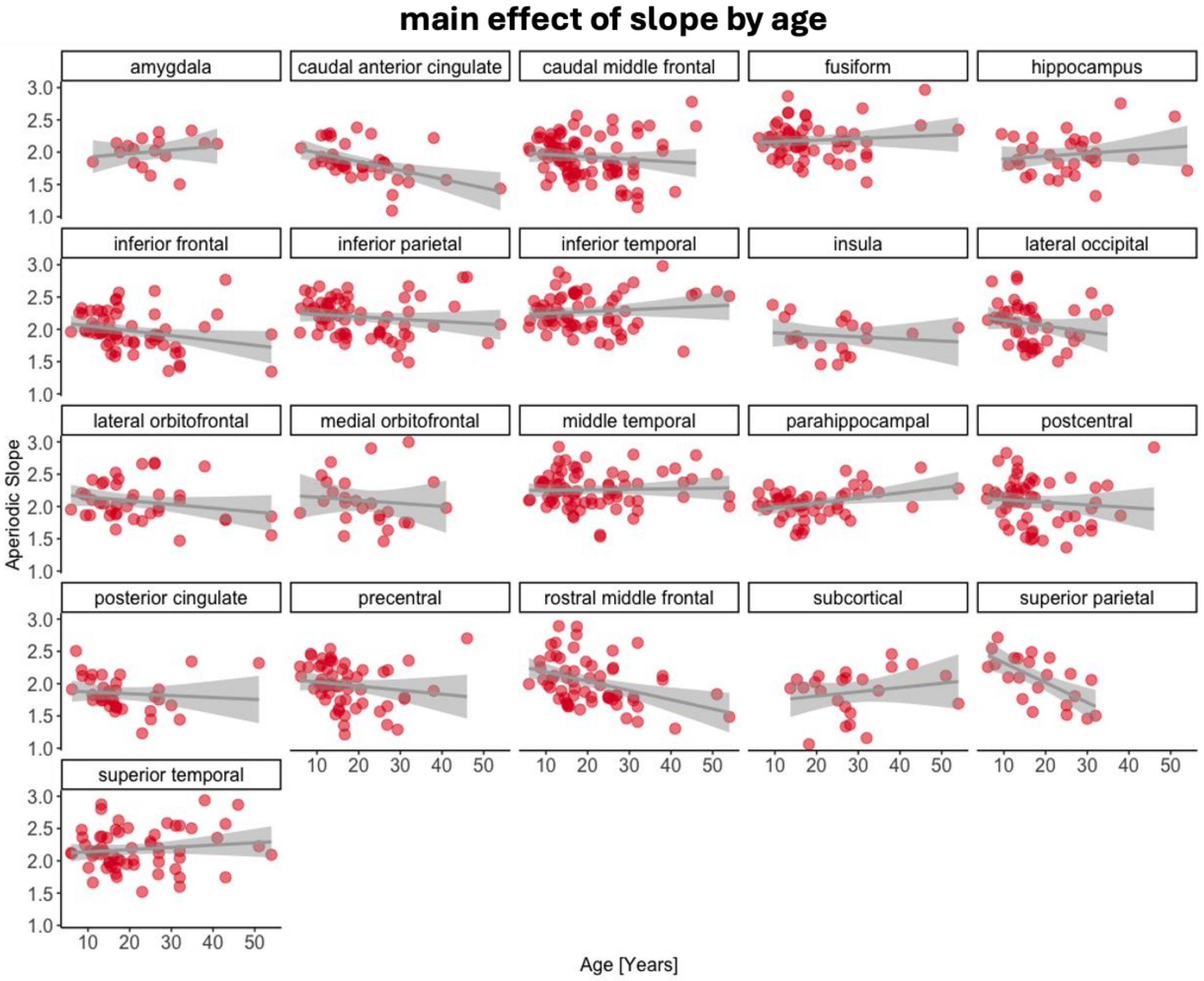


**Figure S4. Relationship between the aperiodic slope and age.** Regions with a statistically significant main effect of age ($p$ <.05) from the linear mixed-effects regression are indicated by the solid red border. The aperiodic slope is on the y-axis, with higher values denoting a steeper slope. Age is on the x-axis, with higher values denoting older age (in years). Data points indicate individual subjects, collapsed across channel and condition (task-based, task-free).

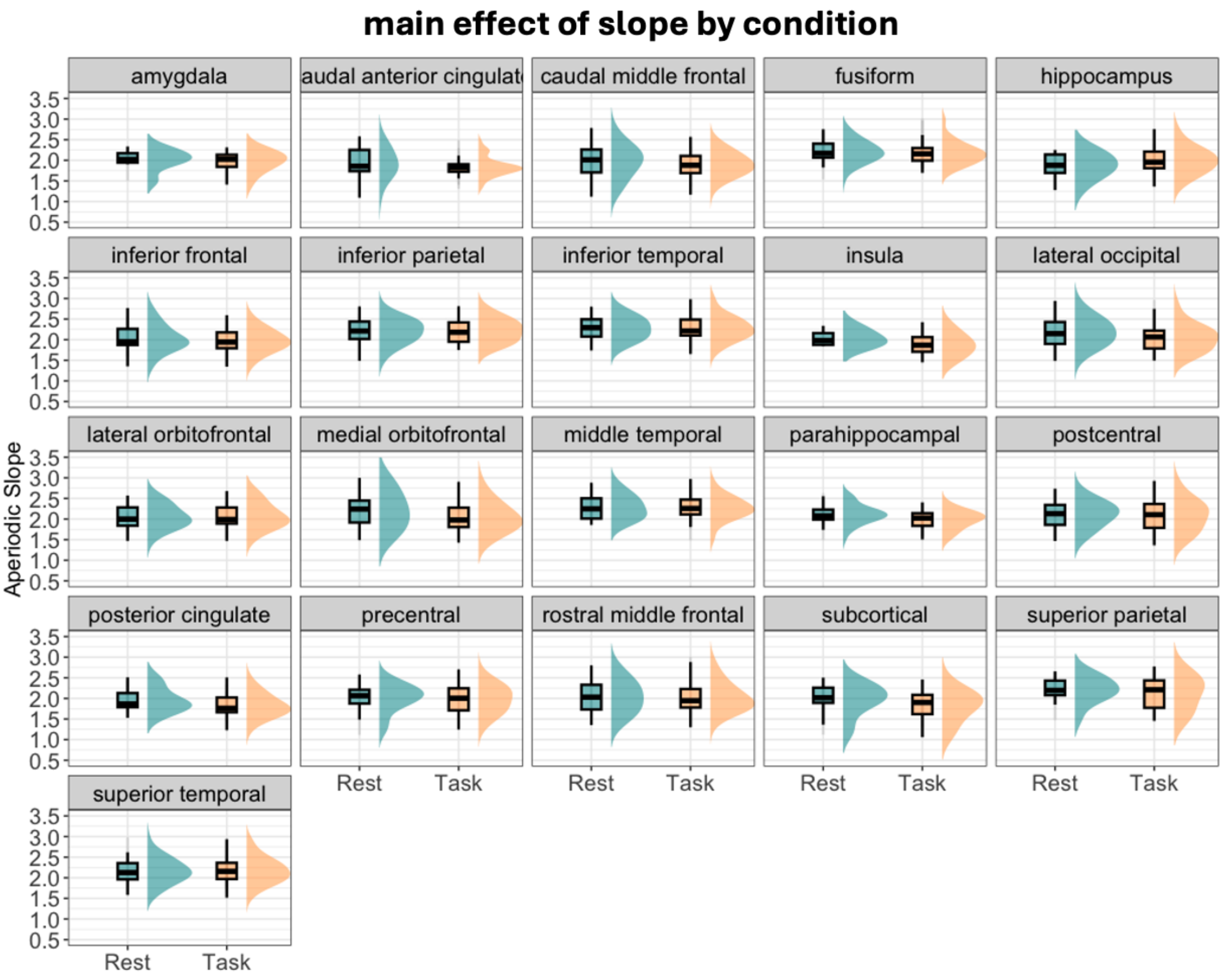


**Figure S5. Differences in the aperiodic slope between conditions (task-based, task-free/rest).** Regions with a statistically significant main effect of condition ($p$ <.05) from the linear mixed-effects regression are indicated by the solid red border. The aperiodic slope is on the y-axis, with higher values denoting a steeper slope. Condition is on the x-axis.

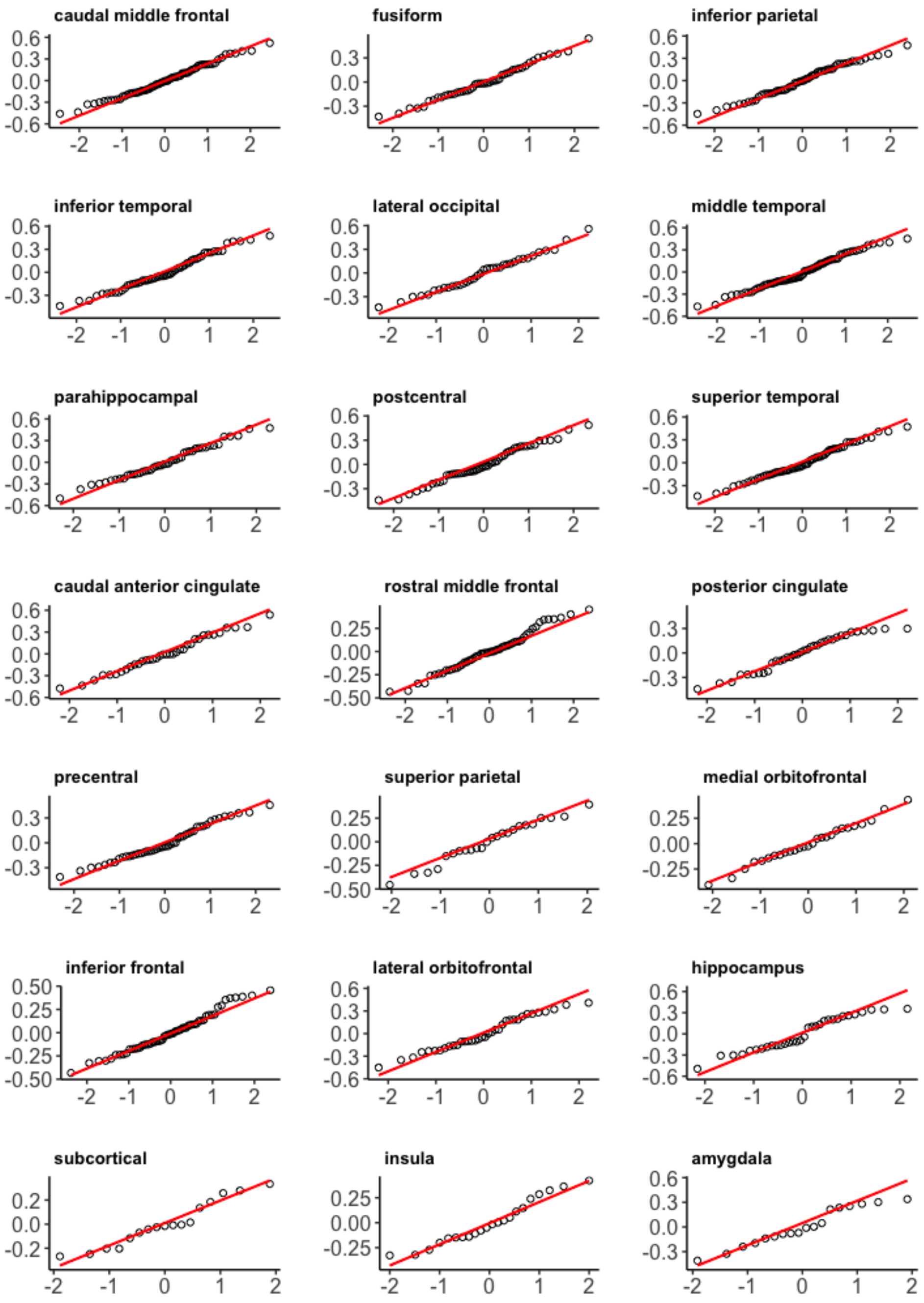


**Figure S6.** Q-Q Plots of residuals for models examining age and aperiodic slopes on memory for each ROI.

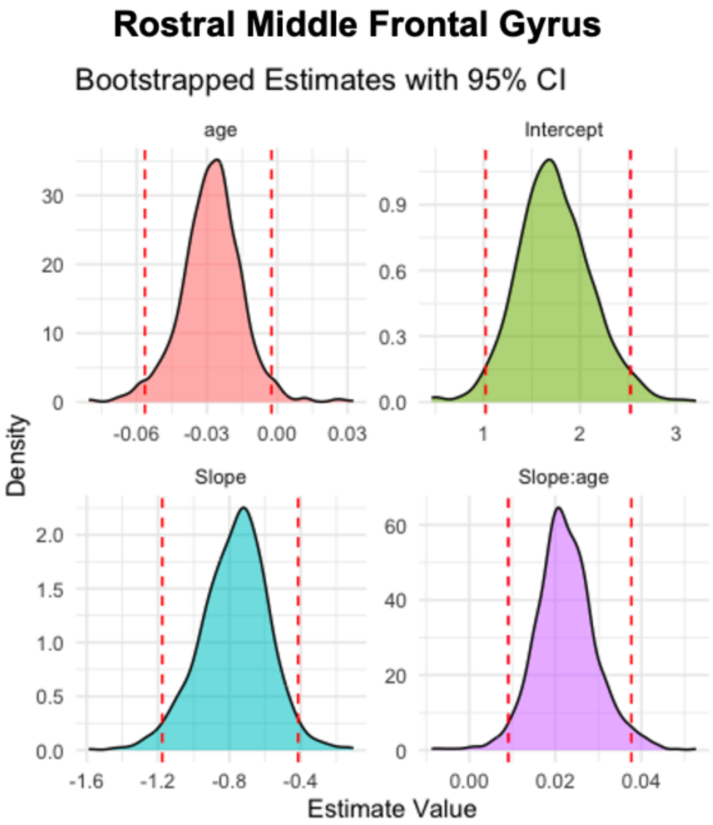


**Figure S7. Bootstrapped beta coefficients for age, slope and memory model for MFG.** Bootstrapped confidence intervals for the intercept, main effects and interactions. Y-axis represents the density of the bootstrapped coefficients and the x-axis represents the estimated beta values. The dashed red lines indicate the 95% confidence interval from the original mixed-model estimates.

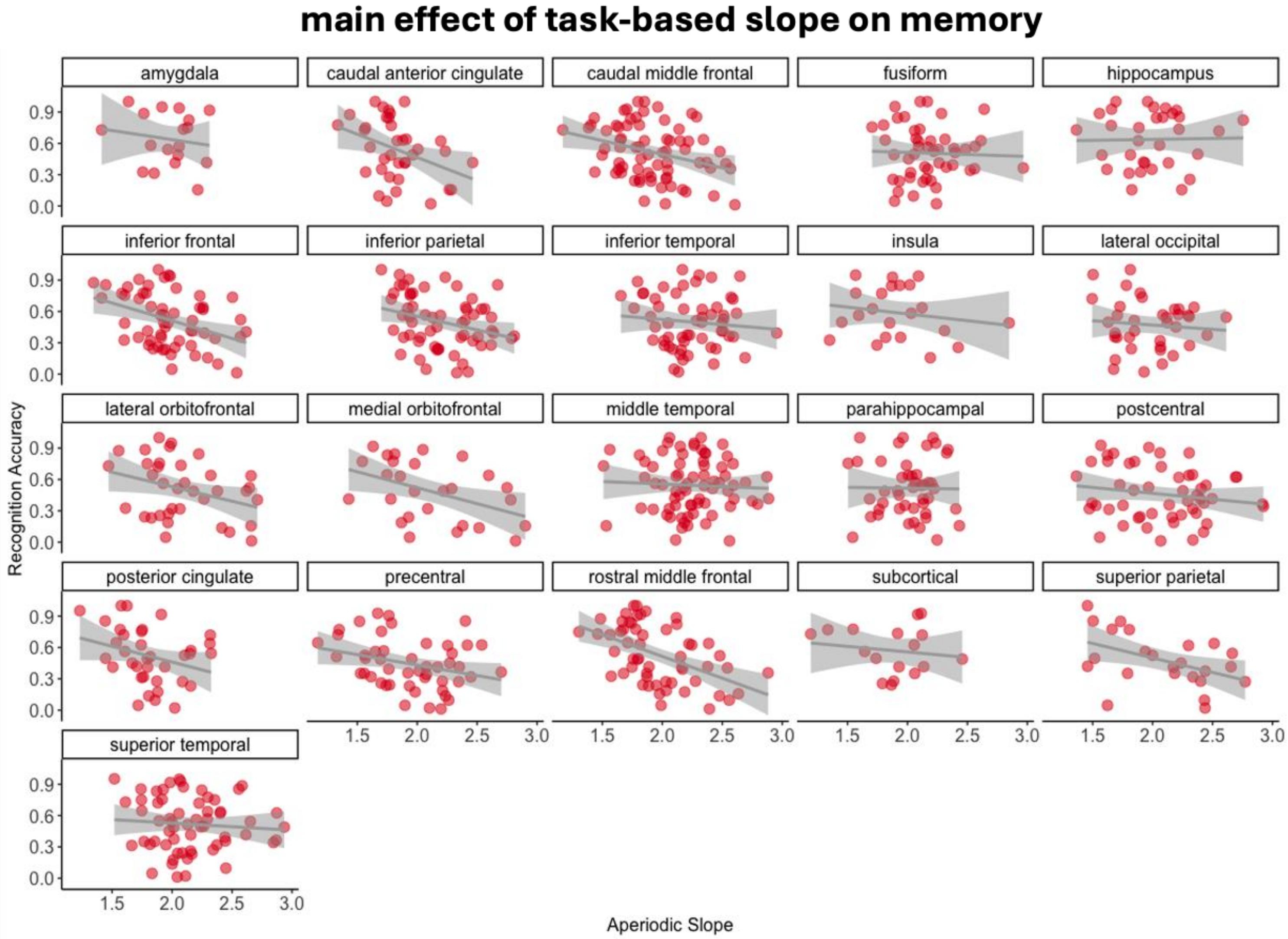


**Figure S8. Relationship between task-based aperiodic slopes and memory performance.** Regions with a statistically significant main effect of slope ($p$ <.05) from the linear mixed-effects regression are indicated by the solid red border. The aperiodic slope is on the x-axis, with higher values denoting a steeper slope. Recognition accuracy is on the y-axis, with higher values denoting better performance. Data points indicate individual subjects, collapsed across channel.

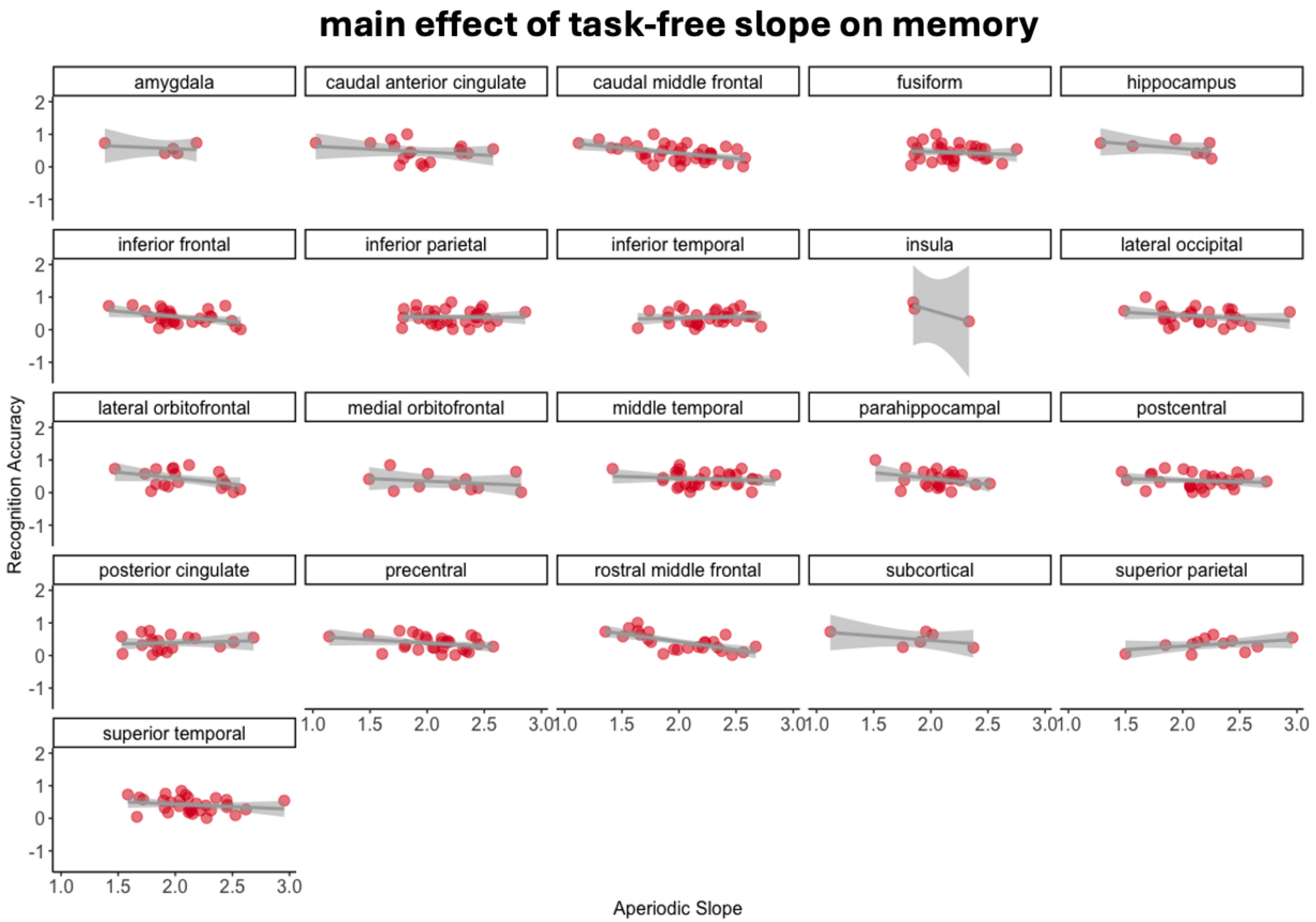


**Figure S9. Relationship between task-free aperiodic slopes and memory performance.** Regions with a statistically significant main effect of slope ($p$ <.05) from the linear mixed-effects regression are indicated by the solid red border. The aperiodic slope is on the x-axis, with higher values denoting a steeper slope. Recognition accuracy is on the y-axis, with higher values denoting better performance. Data points indicate individual subjects, collapsed across channel.

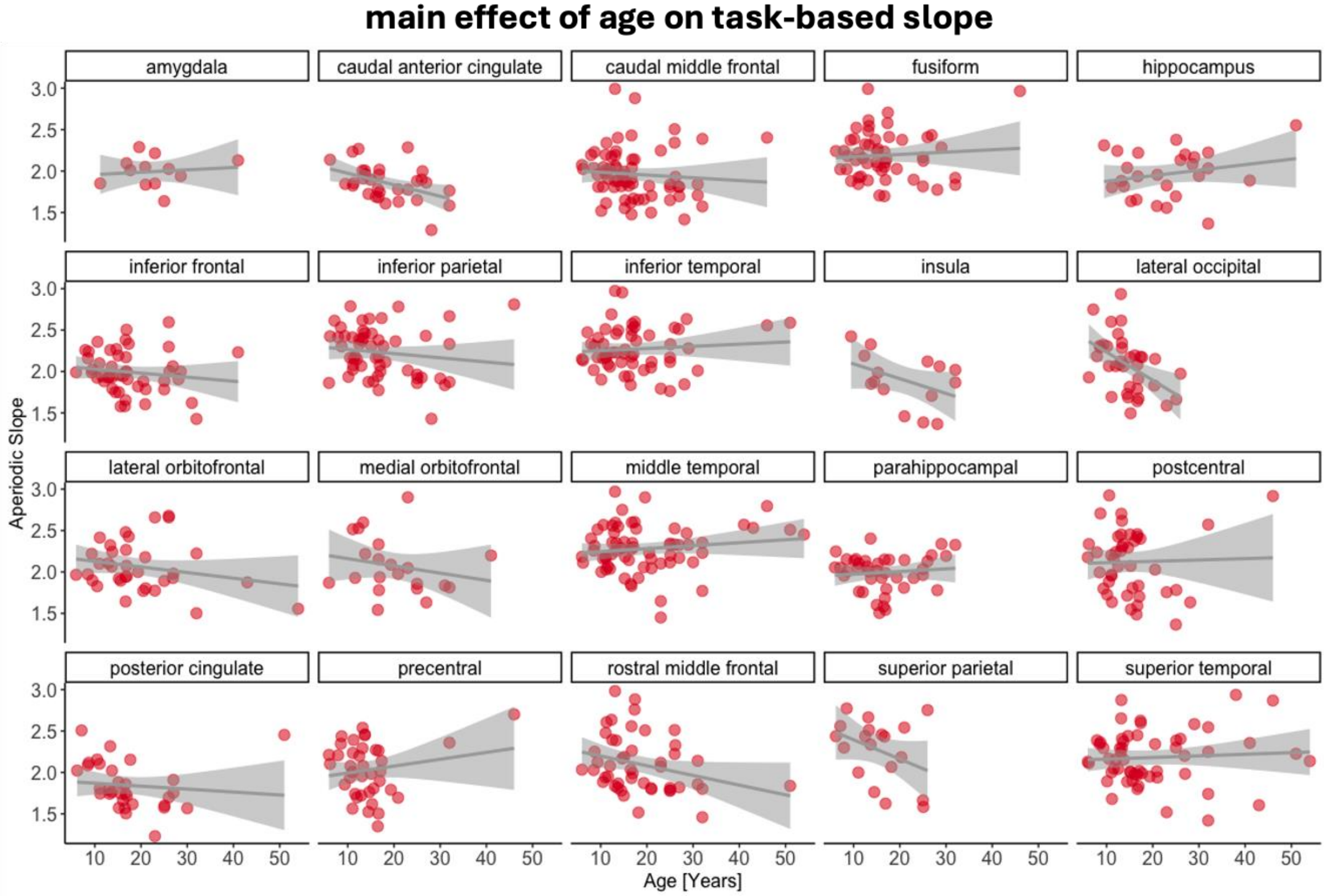


**Figure S10. Relationship between the task-based aperiodic slopes and age from the GMV model.** Regions with a statistically significant main effect of age ($p$ <.05) from the linear mixed-effects regression are indicated by the solid red border. The aperiodic slope is on the y-axis, with higher values denoting a steeper slope. Age is on the x-axis, with higher values denoting older age (in years). Data points indicate individual subjects, collapsed across channel.

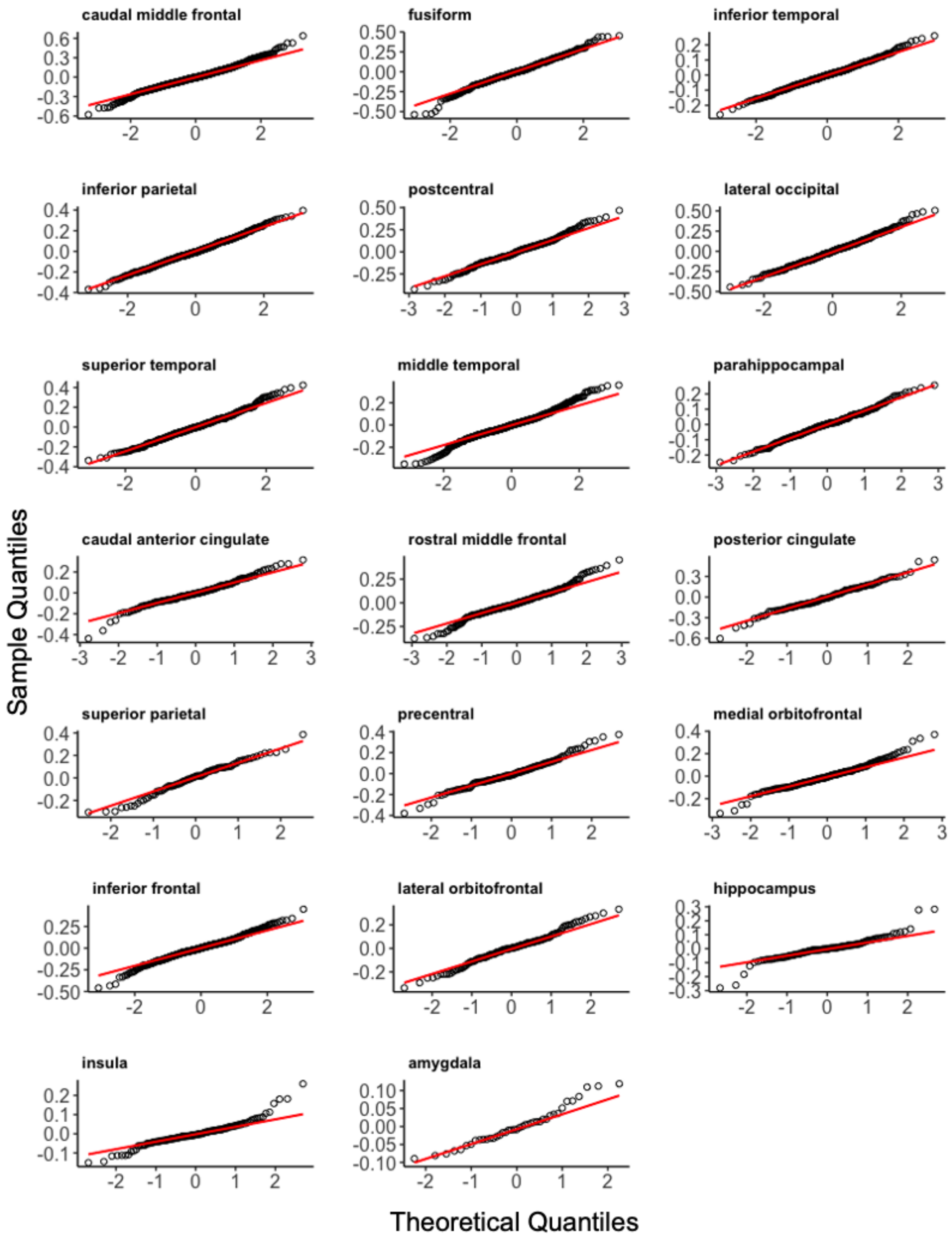


**Figure S11.** Q-Q Plots of residuals for models examining age and GMV on aperiodic slopes for each ROI.

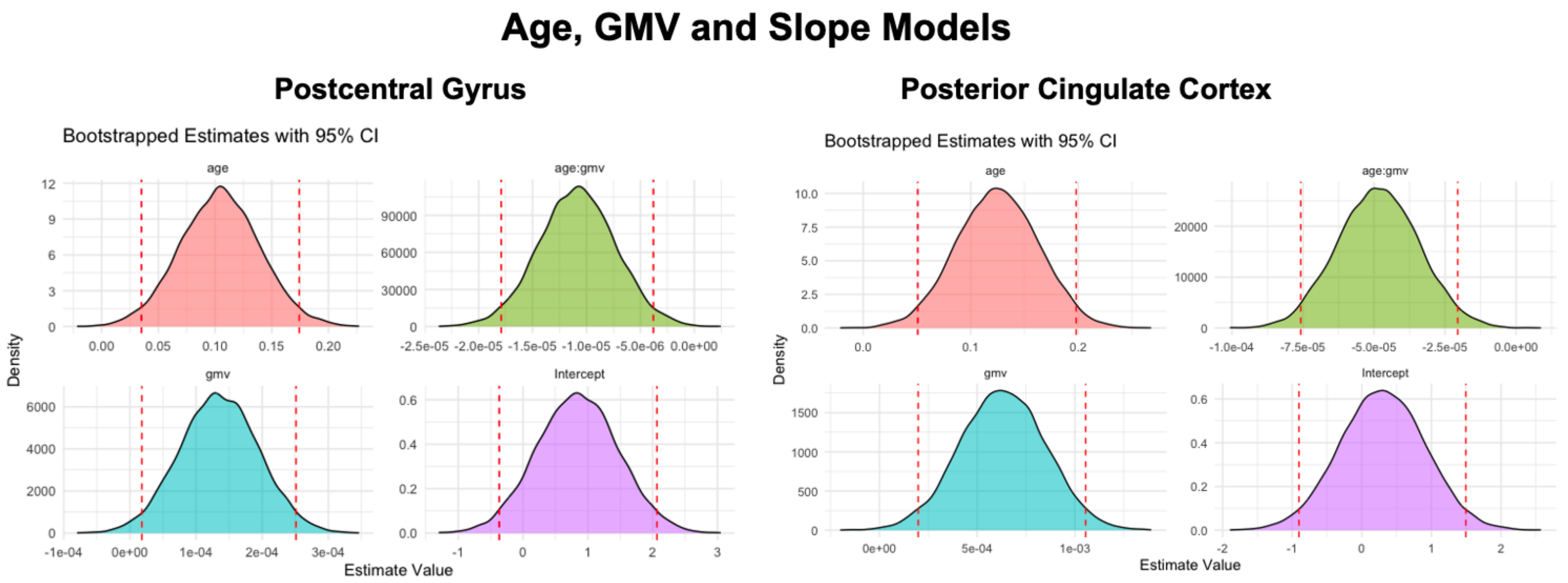


**Figure S12. Bootstrapped beta coefficients for age, GMV and slope models.** Bootstrapped confidence intervals for the intercept, main effects and interactions. Y-axis represents the density of the bootstrapped coefficients and the x-axis represents the estimated beta values. The dashed red lines indicate the 95% confidence interval from the original mixed-model estimates.

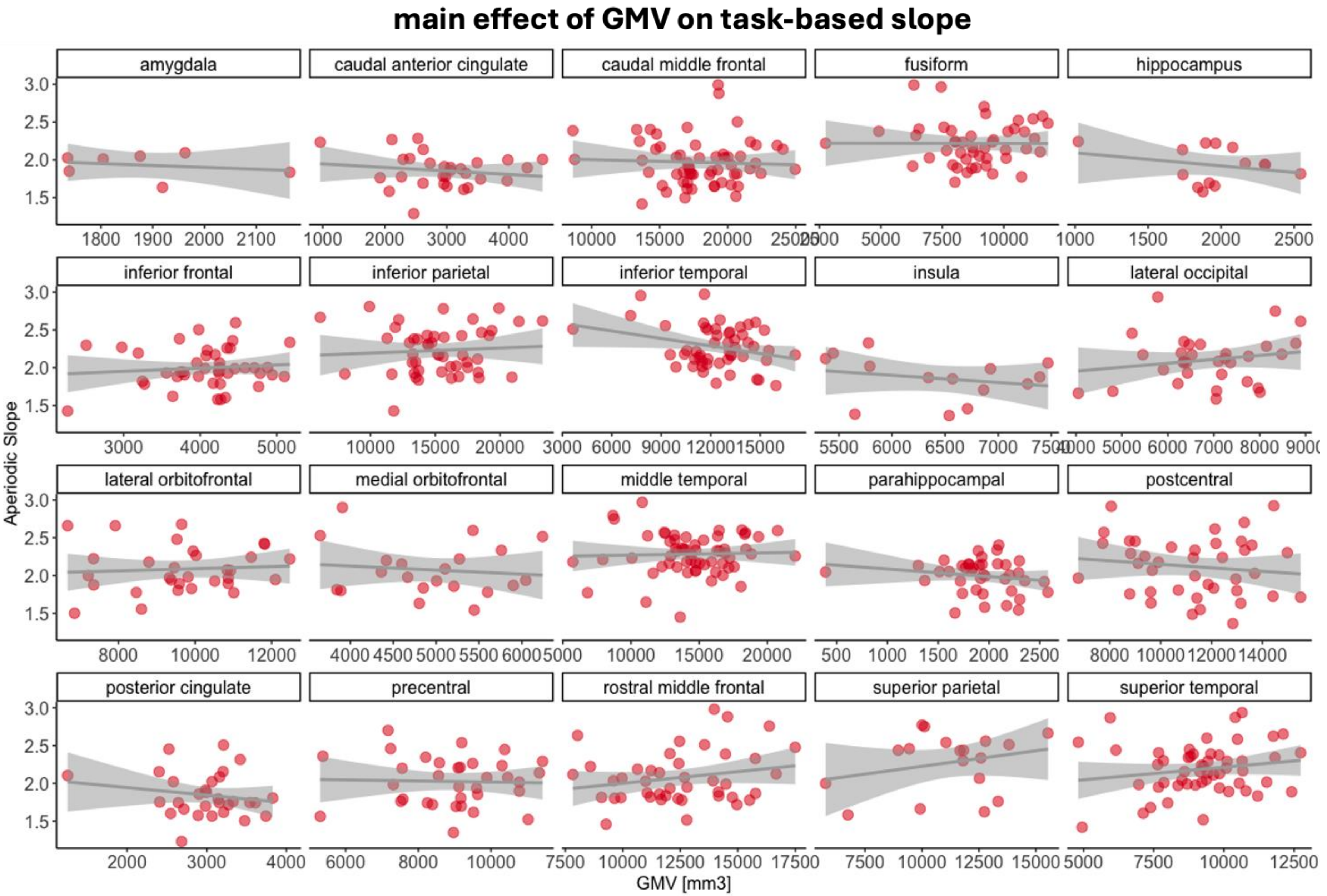


**Figure S13. Relationship between task-based aperiodic slopes and GMV.** Regions with a statistically significant main effect of GMV ($p$ <.05) from the linear mixed-effects regression are indicated by the solid red border. The aperiodic slope is on the y-axis, with higher values denoting a steeper slope. GMV is on the x-axis, with higher values denoting higher GMV. Data points indicate individual subjects, collapsed across channel.

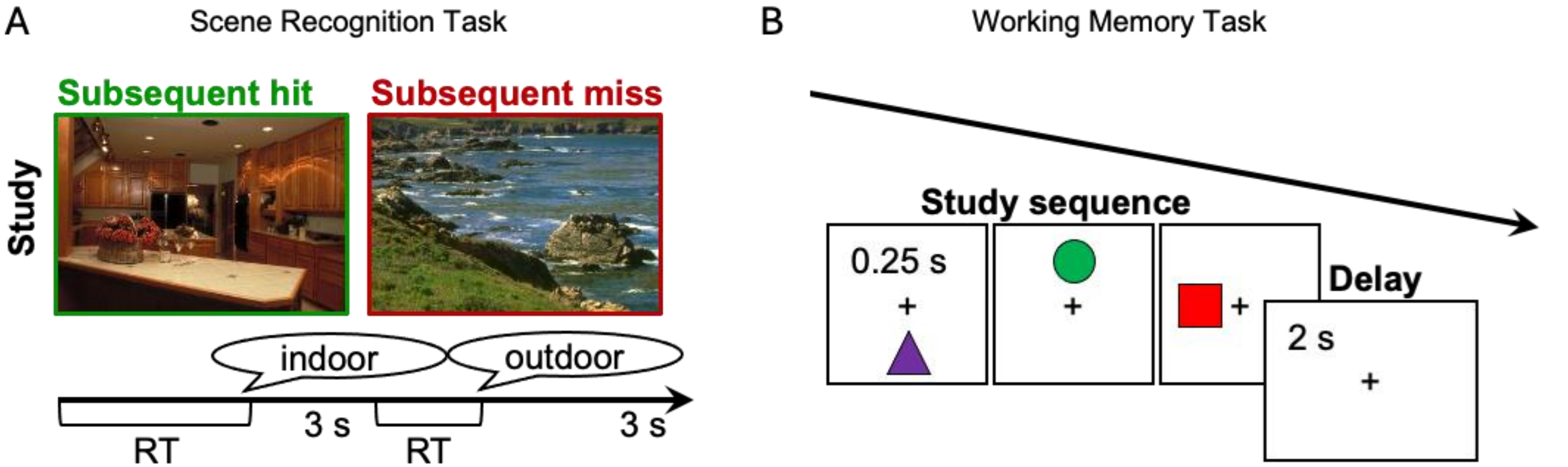


**Figure S14. Schematic of the two visual recognition memory tasks. (A).** Illustration of the visual scene recognition task. Subjects studied sets of 40 pictures of scenes (3s each, separated by a 500ms interstimulus fixation) and made an indoor/outdoor judgment of each scene in preparation for a recognition memory test of all scenes presented during the study block, intermixed with 20 new scenes. **(B)** Illustration of the visual working memory task. Subjects encoded three shapes in a specific spatiotemporal sequence in preparation for a self-paced old/new recognition test of sequences that match exactly or mismatch on one dimension (i.e., shape identity, spatial position, or temporal order).

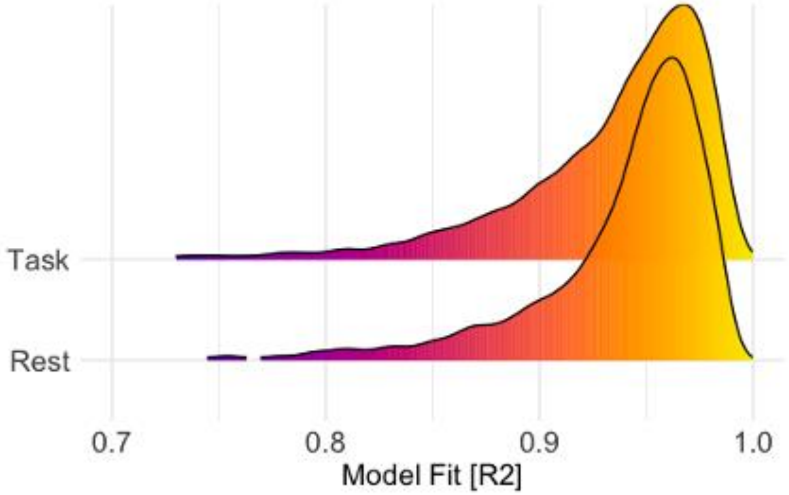


**Figure S15.** Density plots illustrating the distribution of the goodness of fit of (R2) aperiodic model estimation.

**Table S2.** Summary of key dependent and independent variables across each region of interest for task-based recordings.

| Region | Condition | Age Range | | GMV Range | | Memory Range | | Slope Range | | Offset Range | | n Subject | n Channel |
|---|---|---|---|---|---|---|---|---|---|---|---|---|---|
| amygdala | Task | 11.16 | 41 | 1735.5 | 2164.75 | 0.15 | 1 | 1.58 | 2.40 | 3.57 | 6.99 | 14 | 37 |
| caudal anterior cingulate cortex | Task | 6.16 | 32 | 959.25 | 4544.5 | 0.02 | 1 | 1.28 | 2.56 | 1.33 | 9.67 | 31 | 129 |
| caudal middle frontal gyrus | Task | 5.93 | 46 | 8626.25 | 24964.25 | 0.01 | 1 | 1.12 | 3.01 | 1.58 | 11.72 | 68 | 655 |
| fusiform gyrus | Task | 6.16 | 46 | 2752.5 | 11793.5 | 0.02 | 1 | 1.44 | 3.06 | 3.38 | 11.52 | 53 | 249 |
| hippocampus | Task | 9.45 | 51 | 1022.25 | 2544 | 0.15 | 1 | 1.25 | 2.86 | 2.72 | 9.58 | 27 | 108 |
| inferior frontal gyrus | Task | 5.93 | 41 | 2268.83 | 5171.5 | 0.01 | 1 | 1.22 | 2.95 | 1.39 | 11.54 | 51 | 309 |
| inferior parietal cortex | Task | 5.93 | 46 | 6130 | 23277 | 0.01 | 1 | 1.40 | 3.09 | 3.30 | 12.37 | 57 | 395 |
| inferior temporal cortex | Task | 5.93 | 51 | 3638.5 | 17129.5 | 0.02 | 1 | 1.63 | 3.15 | 2.75 | 12.73 | 57 | 240 |
| insula | Task | 9.45 | 38 | 5374.5 | 7468.5 | 0.15 | 0.94 | 1.16 | 2.71 | 3.92 | 8.23 | 16 | 92 |
| lateral occipital cortex | Task | 6.16 | 26 | 4049.75 | 8889.12 | 0.02 | 1 | 1.18 | 3.02 | 2.30 | 13.37 | 38 | 213 |
| lateral orbitofrontal cortex | Task | 5.93 | 54 | 6668 | 12474.5 | 0.01 | 1 | 1.27 | 2.90 | 1.86 | 11.49 | 36 | 96 |
| medial orbitofrontal cortex | Task | 5.93 | 41 | 3649 | 6241.5 | 0.01 | 0.91 | 1.29 | 3.17 | 3.66 | 11.53 | 23 | 128 |
| middle temporal cortex | Task | 5.93 | 54 | 5741.5 | 22048 | 0.01 | 1 | 1.45 | 3.15 | 1.48 | 11.68 | 68 | 411 |
| parahippocampal gyrus | Task | 5.93 | 38 | 392.25 | 2579.75 | 0.02 | 1 | 1.39 | 2.64 | 3.75 | 10.32 | 44 | 144 |
| postcentral gyrus | Task | 5.93 | 46 | 6759.5 | 15504.5 | 0.01 | 0.92 | 1.33 | 3.06 | 3.12 | 11.32 | 45 | 149 |
| posterior cingulate cortex | Task | 6.16 | 51 | 1252 | 4043 | 0.02 | 1 | 1.23 | 2.68 | 1.24 | 11.67 | 34 | 87 |
| precentral gyrus | Task | 5.93 | 46 | 5302.25 | 11415 | 0.01 | 0.92 | 1.14 | 2.98 | 3.58 | 11.05 | 38 | 85 |
| rostral middle frontal gyrus | Task | 5.93 | 51 | 7832 | 17498 | 0.01 | 1 | 1.28 | 3.06 | 2.19 | 11.50 | 47 | 188 |
| subcortical | Task | 9.45 | 51 | NA | NA | 0.24 | .92 | 1.13 | 2.31 | 1.50 | 8.32 | 16 | 56 |
| superior parietal cortex | Task | 6.16 | 26 | 5766.66 | 15529.16 | 0.02 | 0.85 | 1.46 | 2.93 | 2.90 | 12.59 | 20 | 60 |
| superior temporal cortex | Task | 5.93 | 54 | 4813.25 | 12732.75 | 0.01 | 0.92 | 1.23 | 3.08 | 1.18 | 10.94 | 58 | 248 |

**Table S3.** Summary of key dependent and independent variables across each region of interest for task-free recordings.

| Region | Condition | Age Range | | GMV Range | | Slope Range | | Offset Range | | n Subject | n Channel |
|---|---|---|---|---|---|---|---|---|---|---|---|
| amygdala | Rest | 16.84 | 34.84 | 1296.25 | 1962.50 | 1.85 | 2.34 | 3.80 | 6.15 | 7 | 13 |
| caudal anterior cingulate cortex | Rest | 6.17 | 32.00 | 959.25 | 4544.50 | 1.32 | 2.62 | 2.23 | 8.82 | 18 | 61 |
| caudal middle frontal gyrus | Rest | 5.94 | 45.00 | 8753.50 | 24964.25 | 1.11 | 3.03 | 2.80 | 11.19 | 43 | 364 |
| fusiform gyrus | Rest | 6.17 | 54.00 | 2752.50 | 11793.50 | 1.54 | 3.11 | 3.55 | 11.44 | 44 | 242 |
| hippocampus | Rest | 9.46 | 54.00 | 1482.75 | 2327.25 | 1.20 | 2.70 | 2.11 | 9.34 | 13 | 49 |
| inferior frontal gyrus | Rest | 5.94 | 54.00 | 2268.83 | 5100.00 | 1.22 | 2.97 | 1.62 | 11.93 | 34 | 208 |
| inferior parietal cortex | Rest | 5.94 | 54.00 | 10405.00 | 23277.00 | 1.41 | 3.14 | 4.46 | 10.02 | 41 | 219 |
| inferior temporal cortex | Rest | 5.94 | 54.00 | 3638.50 | 17129.50 | 1.60 | 3.16 | 2.52 | 10.72 | 41 | 184 |
| insula | Rest | 9.46 | 54.00 | 4852.00 | 6928.00 | 1.39 | 2.62 | 3.69 | 8.09 | 7 | 31 |
| lateral occipital cortex | Rest | 6.17 | 34.84 | 4801.50 | 8889.13 | 1.21 | 3.10 | 3.27 | 9.66 | 33 | 143 |
| lateral orbitofrontal cortex | Rest | 5.94 | 54.00 | 6849.00 | 12474.50 | 1.24 | 2.80 | 2.94 | 11.67 | 25 | 62 |
| medial orbitofrontal cortex | Rest | 5.94 | 32.00 | 3843.00 | 6241.50 | 1.56 | 3.12 | 3.29 | 11.56 | 12 | 37 |
| middle temporal cortex | Rest | 5.94 | 54.00 | 5741.50 | 22048.00 | 1.53 | 3.11 | 2.98 | 10.98 | 41 | 228 |
| parahippocampal gyrus | Rest | 5.94 | 54.00 | 392.25 | 2352.50 | 1.38 | 2.81 | 3.73 | 9.50 | 33 | 120 |
| postcentral gyrus | Rest | 5.94 | 20.50 | 6759.50 | 14982.50 | 1.37 | 2.87 | 4.37 | 9.41 | 29 | 90 |
| posterior cingulate cortex | Rest | 6.17 | 43.00 | 1252.00 | 4043.00 | 1.50 | 2.66 | 3.08 | 9.17 | 21 | 50 |
| precentral gyrus | Rest | 5.94 | 20.50 | 5302.25 | 11415.00 | 1.28 | 2.79 | 4.12 | 10.07 | 26 | 51 |
| rostral middle frontal gyrus | Rest | 5.94 | 32.00 | 9210.50 | 17498.00 | 1.24 | 3.04 | 2.75 | 10.68 | 25 | 93 |
| subcortical | Rest | 9.46 | 54.00 | NA | NA | 1.10 | 2.62 | 1.18 | 7.77 | 13 | 49 |
| superior parietal cortex | Rest | 6.17 | 43.00 | 5766.67 | 15529.17 | 1.50 | 2.99 | 3.91 | 9.04 | 15 | 30 |
| superior temporal cortex | Rest | 5.94 | 43.00 | 4951.50 | 12732.75 | 1.33 | 3.07 | 3.27 | 9.98 | 36 | 164 |

**Table S4.** Sex, age, seizure onset zone and iEEG recording type for each patient.

| Subject | Sex | Age | Seizure Onset Zone | Type |
|---|---|---|---|---|
| S1 | F | 5.94 | R F | ECoG |
| S2 | F | 6.17 | L PO | ECoG |
| S3 | M | 7.12 | R TP | sEEG |
| S4 | F | 7.92 | NC | ECoG |
| S5 | M | 8.5 | L P | combined |
| S6 | F | 8.67 | L P | combined |
| S7 | M | 9.33 | L T | ECoG |
| S8 | M | 9.46 | R T | sEEG |
| S9 | F | 10.13 | UN | ECoG |
| S10 | F | 10.5 | R T | ECoG |
| S11 | M | 10.58 | R P | ECoG |
| S12 | M | 11 | L T | ECoG |
| S13 | M | 11.03 | UN | sEEG |
| S14 | M | 11.08 | B F | ECoG |
| S15 | M | 11.17 | UN | sEEG |
| S16 | M | 11.25 | L OT | ECoG |
| S17 | F | 11.58 | NC | ECoG |
| S18 | F | 12 | L FP | ECoG |
| S19 | M | 12.33 | UN | sEEG |
| S20 | F | 12.5 | R P | ECoG |
| S21 | F | 12.5 | L F | combined |
| S22 | M | 12.92 | L T | ECoG |
| S23 | F | 13.08 | R TPO | ECoG |
| S24 | M | 13.1 | L FT | sEEG |
| S25 | M | 13.17 | L O | combined |
| S26 | F | 13.21 | L insula | sEEG |
| S27 | M | 13.33 | R F | combined |
| S28 | M | 13.33 | R F | ECoG |
| S29 | F | 13.67 | L T | ECoG |
| S30 | F | 13.7 | R P | sEEG |
| S31 | M | 13.77 | UN | sEEG |

| | | | | |
|---|---|---|---|---|
| S32 | M | 14.46 | L T | sEEG |
| S33 | F | 14.67 | R T | combined |
| S34 | M | 15 | UN | sEEG |
| S35 | M | 15.08 | R F | ECoG |
| S36 | F | 15.21 | R TPO | ECoG |
| S37 | M | 15.27 | L insula | sEEG |
| S38 | M | 15.58 | L T | ECoG |
| S39 | F | 16.08 | R P | combined |
| S40 | M | 16.5 | R F | sEEG |
| S41 | M | 16.67 | R FT | ECoG |
| S42 | F | 16.67 | NC | ECoG |
| S43 | M | 16.67 | NC | ECoG |
| S44 | F | 16.75 | R F | sEEG |
| S45 | F | 16.84 | R T | sEEG |
| S46 | M | 16.92 | NC | ECoG |
| S47 | M | 17.08 | NC | combined |
| S48 | M | 17.33 | L T | ECoG |
| S49 | M | 17.42 | L F | combined |
| S50 | M | 17.69 | UN | sEEG |
| S51 | M | 18.17 | B P | sEEG |
| S52 | F | 19.33 | L F | ECoG |
| S53 | M | 19.6 | L T | sEEG |
| S54 | F | 20.33 | L O | sEEG |
| S55 | M | 20.5 | R TF | ECoG |
| S56 | F | 20.9 | UN | sEEG |
| S57 | F | 20.97 | B generalized | sEEG |
| S58 | F | 21 | B T | sEEG |
| S59 | M | 23 | UN | combined |
| S60 | M | 23 | UN | sEEG |
| S61 | M | 23 | L TP | combined |
| S62 | M | 25 | R T, O, R F | ECoG |
| S63 | F | 25 | R SMA | sEEG |

| | | | | |
|---|---|---|---|---|
| S64 | M | 25 | UN | ECoG |
| S65 | F | 25.08 | B multifocal | sEEG |
| S66 | M | 26 | NC | ECoG |
| S67 | M | 26 | BT | sEEG |
| S68 | M | 26 | R T | sEEG |
| S69 | M | 26.15 | R FT | sEEG |
| S70 | M | 26.84 | L T | sEEG |
| S71 | M | 26.89 | L FT | sEEG |
| S72 | M | 27 | R FT | sEEG |
| S73 | M | 27 | B T | sEEG |
| S74 | F | 27 | L anterior insula | sEEG |
| S75 | F | 28 | R P | sEEG |
| S76 | F | 28.11 | R T | sEEG |
| S77 | F | 28.59 | L T | sEEG |
| S78 | M | 29 | L T | sEEG |
| S79 | F | 29.34 | R motor | sEEG |
| S80 | M | 30 | UN | sEEG |
| S81 | M | 31 | NC | ECoG |
| S82 | M | 31 | UN | ECoG |
| S83 | F | 31 | R T | sEEG |
| S84 | M | 32 | UN | ECoG |
| S85 | F | 32 | L UN | sEEG |
| S86 | M | 32 | UN | sEEG |
| S87 | M | 32 | R T | sEEG |
| S88 | F | 32 | B FT | sEEG |
| S89 | M | 32 | L hippocampus/amygdala/insula | sEEG |
| S90 | M | 34.84 | R TP | sEEG |
| S91 | M | 38 | L T | sEEG |
| S92 | F | 38 | UN | sEEG |
| S93 | M | 38 | L F | sEEG |
| S94 | M | 41 | B T | sEEG |
| S95 | M | 43 | B T | sEEG |

| | | | | |
|---|---|---|---|---|
| S96 | F | 43 | UN | sEEG |
| S97 | M | 45 | L hippocampus | ECoG |
| S98 | M | 46 | UN | ECoG |
| S99 | F | 51 | UN | sEEG |
| S100 | M | 54 | UN | sEEG |
| S101 | M | 54 | R T | sEEG |

*Note.* L = left; R = right; B = bilateral; F = frontal lobe; T = temporal lobe; P = parietal lobe; O = occipital lobe; NC = seizures not captured; UN = unknown.
